# NIRSTORM: a Brainstorm extension dedicated to functional Near Infrared Spectroscopy (fNIRS) data analysis, advanced 3D reconstructions, and optimal probe design

**DOI:** 10.1101/2024.09.05.611463

**Authors:** Édouard Delaire, Thomas Vincent, Zhengchen Cai, Alexis Machado, Laurent Hugueville, Denis Schwartz, Francois Tadel, Raymundo Cassani, Louis Bherer, Jean-Marc Lina, Mélanie Pélégrini-Issac, Christophe Grova

## Abstract

**Significance:** We propose NIRSTORM, a software package built within Brainstorm environment, enabling full data analysis of functional Near InfraRed Spectroscopy (fNIRS) data from experiment planning to 3D reconstruction of hemodynamic fluctuations on the cortical surface using optical tomographic approaches. NIRSTORM enables the integration of fNIRS analysis within a multimodal setup making it easy to study fNIRS in combination with other multimodal data such as electroencephalography (EEG) or magnetic resonance imaging (MRI).

**Aim:** NIRSTORM aims to provide an easy-to-use and fully modular toolbox for fNIRS analysis from experimental planning to optical tomography 3D reconstruction extending Brainstorm capacity for multimodal analysis.

**Approach:** NIRSTORM was developed in MATLAB® and integrated as a plugin of the software Brainstorm. Brainstorm is a GUI-oriented, widely used software originally dedicated to statistical analysis and source imaging of EEG and magnetoencephalography (MEG) data.

**Results:** In addition to conventional fNIRS preprocessing steps, including standard channel space and statistical analyses, NIRSTORM provides advanced methods dedicated to optimal probe placement, allowing personalized fNIRS study designs and accurate near-infrared optical tomography within the Maximum Entropy on the Mean (MEM) framework.

**Conclusion:** NIRSTORM is an open-access, user-friendly plugin extending the capacity of Brainstorm, for fNIRS analysis, therefore narrowing the gap between EEG/MEG and hemodynamics.

## 1. Introduction

Functional near-infrared spectroscopy (fNIRS) is a wearable, noninvasive imaging technique able to monitor brain hemodynamic activity^1–4^. Using sensors attached to the head and without requiring strict immobility like other neuroimaging techniques (functional magnetic resonance imaging – fMRI, Magnetoencephalography – MEG), fNIRS is an ideal technique to monitor brain activity in experimental designs mimicking realistic lifestyle conditions (e.g., while walking^5^, or sleeping^6^). fNIRS is also well adapted for prolonged recordings, in clinical settings notably (e.g., epilepsy^7^, intensive care units^8^). fNIRS consists of installing a set of optodes (light emitting sources, and light detectors) on the scalp of a participant. Light sources emit near-infrared light using two or more wavelengths typically in the range of 650 to 900 nm. Within this optical window, since water is not absorbing much light, NIR light can travel long distances in living tissues with little absorption but large scattering. From the light intensity measured on fNIRS detectors, one can then estimate, with good temporal accuracy, the local variations in oxyhemoglobin (HbO), deoxyhemoglobin (HbR), and total hemoglobin (HbT = HbO + HbR) concentrations between a source and a detector (also called a fNIRS “channel”) using at least two wavelengths^4^. Similarly to functional magnetic resonance imaging (fMRI), fNIRS is an indirect measure of brain activity, in that it consists in monitoring the hemodynamic processes elicited by neuronal discharges, thanks to the so-called neurovascular coupling mechanism^9^. However, as opposed to fMRI, which allows whole-brain measurements, fNIRS is only sensitive to the superficial cortical layer, due to the nature of near-infrared light scattering and absorption and the use of sensors attached to the scalp ^10,11^.

A standard fNIRS experiment can be divided into three steps, as described in further detail in Yücel et al, 2021^12^: acquisition planning, data acquisition, and data analysis.

Regarding acquisition planning, the main objective of those algorithms is to decide, given a targeted region of interest (ROI) in the brain and a set of additional constraints on the montage (e.g. the number of available optodes), where to place the sources and detectors while maximizing the sensitivity of the montage to the targeted ROIs. Three main methods have been proposed to assist the placement of fNIRS sensors for a specific experiment: fNIRS optodes’ location decider^13^, personalized optimal montage (developed by our team and implemented in NIRSTORM)^14,15^ and Array designer ^16^. “fNIRS optodes’ location decider” assigns the location of the optodes from a set of predefined positions from the standard 10-10 or 10-05 international EEG system while providing NIR light sensitivity maps estimated on two head templates (Colin27^17^, ICBM152^18^). The main limitation of this approach is that the anatomical information of the template might not accurately reflect the underlying anatomy of the studied population. Additionally, the set of proposed positions coming from the 10-10 system might not allow the realization of an fNIRS montage spatially dense enough to enable accurate 3D tomographic reconstruction of the fluctuations of HbO/HbR in the brain. To solve this issue, we proposed the concept of a personalized optimal montage algorithm allowing using the subject-specific anatomical information to optimize the fNIRS montage ^14,15^. A similar approach is also implemented in Array Designer^16^. In the implementation made in NIRSTORM, we included a constraint on the montage to favor locally dense montage enabling locally accurate reconstruction of the concentration changes on the cortical surface, while keeping the possibility to add EEG electrodes following a standard EEG montage.

In the context of fNIRS data analysis, several toolboxes have been proposed with many teams also using their in-house code. An exhaustive list of the existing toolboxes and their comparison is out of the scope of this current article (see https://fnirs.org/resources/data-analysis/software/ for a list of software). Among those toolboxes, a few of them are based, like ours, on existing toolboxes dedicated to EEG/MEG analysis such as MNE python^19^ and Fieldtrip^20^. These two toolboxes focused on the analysis of fNIRS signals at the sensor level, including the possibility of performing advanced statistical analysis using a general linear model (GLM) to estimate brain activation. On the other hand, few other toolboxes decided to leverage the similarity of fNIRS signal analysis with conventional fMRI analysis packages, by creating plugins for well-established toolboxes such as SPM^21^: for example, NIRS-SPM^22^, LIONirs^23^, or NIRS-Kit^24^. Finally, a few other toolboxes offer the possibility to apply fNIRS tomographic reconstruction to estimate the hemodynamic response along the cortical surface or in the brain, using mainly variants of the minimum norm estimate reconstruction method (e.g. NeuroDot^25,26^, the NIRS brain AnalyzIR toolbox^27^, and AtlasViewer^28^).

In this context, we developed NIRSTORM as an open-source toolbox dedicated to fNIRS analysis from experimental planning to optical tomography 3D reconstruction. NIRSTORM is fully integrated within Brainstorm, a widely used open-source software dedicated to the analysis of MEG, EEG, iEEG, and multiunit electrophysiology^29,30^. NIRSTORM thus extends Brainstorm’s features by adding the possibility to process and reconstruct fNIRS data and analyze multimodal signals. Our rationale for incorporating NIRSTORM in Brainstorm as a package dedicated to EEG/MEG analysis is that EEG/MEG and fNIRS signals share many similarities, even if they are measuring signals of different physiological origins: (i) they consist of scalp measurements usually resulting in long duration recording requiring data visualization/interaction, (ii) they offer an excellent temporal resolution, (iii) their spatial resolution remains limited, either as interpretation from scalp recordings or after applying 3D reconstruction of the underlying cortical generators of these scalp recordings, which requires solving an ill-posed inverse problem. Brainstorm software is internationally recognized for EEG/MEG processing^30^ featuring advanced databasing, visualization, signal processing, source localization, and statistical analysis methods. The similarities between EEG, MEG, and fNIRS allow us to leverage tools developed in Brainstorm studying fNIRS.

This article aims to introduce the main features of the open-source NIRSTOM toolbox and is organized as follows: in section 2, we provide the reader with an overview of the toolbox. In section 3, we briefly describe how to design a personalized optimal montage that maximizes the sensitivity of fNIRS recordings to reach specific targeted regions of interest. In section 4, we describe the basic features of channel-space analysis such as data importation, preprocessing, and statistical modeling. Finally, in section 5, we provide further details on 3D Near Infra-red Optical Tomography (NIROT) reconstruction of fNIRS signals, describing first the computation of the forward model and two methods to solve the inverse problem of fNIRS 3D reconstruction, namely the Minimum Norm Estimation (MNE) and Maximum Entropy on the Mean (MEM) methods. Source code and tutorials are freely available on the GitHub (https://github.com/Nirstorm/nirstorm – https://neuroimage.usc.edu/brainstorm/Tutorials).

## 2. Overview of NIRSTORM

Most illustrations provided in this paper are based on an fNIRS dataset acquired on a right-handed healthy control during a finger-tapping task using their left hand. The protocol and the general workflow from fNIRS optimal montage design to 3D reconstruction is described in further detail in Cai et al, 2021^31^.

### 2.1. Rationale for integrating NIRSTORM into Brainstorm

fNIRS shares several similarities with EEG/MEG data. First, both modalities consider sensors placed on the subject’s head. Therefore, it is important to coregister sensor positions with either the subject’s anatomy or a template when no individual MR structural image is available. Brainstorm user interface already provides the user with the functionalities to load, view, and process MR images and coregister the sensor space with anatomical data. Second, visualization tools and pre-processing methods developed for EEG/MEG and available in Brainstorm can easily be adapted to handle fNIRS data. These methods include: event labeling, noisy channel detection and rejection, principal or independent component analysis (PCA/ICA), temporal filtering, time-frequency analysis, and functional connectivity, among others.

Besides, our team has developed several additional processes specific to fNIRS data processing detailed further in sections 3 and 4.

### 2.2. Database organization and available processes

Brainstorm database organization is protocol-oriented and structured hierarchically so that each subject’s folder can contain multiple data (structural, functional) and experimental conditions. Since NIRSTORM is a Brainstorm plugin implemented within the MATLAB® environment, files are stored in the database as MATLAB® structures. All structures are described in Brainstorm tutorials (https://neuroimage.usc.edu/brainstorm/Tutorials). NIRSTORM users thus benefit from Brainstorm interface and can apply processes and pipelines to their data using the “drag and drop” feature or search queries ^29,30^.

Brainstorm databasing system allows storing and organizing within the same environment anatomical data (raw anatomical MRI and segmented surfaces), fNIRS data (fNIRS time series and sensor locations), but also electrophysiology data when available (EEG, MEG, or intracranial EEG time series, together with sensor locations), as well as additional physiological measurements (e.g., cardiac and respiration monitoring, pulse oximetry, accelerometers). Spatial registration is ensured through sensor digitalization and coregistration procedures, while the incorporation of specific triggers associated with the time series allows synchronization in time.

### 2.3. Improving numerical reproducibility

To provide a user-friendly framework to easily reproduce fNIRS analysis pipelines, NIRSTORM processes as well as Brainstorm ones were added within the Brainstorm pipeline scripting system to facilitate file management and allow the user to re-run some parts or the whole fNIRS data analysis pipeline while offering the possibility to modify one or several parameters. Examples of these scripts specifically designed for fNIRS analysis, including the ones used in this manuscript, are available for download on NIRSTORM project’s GitHub page (https://github.com/Nirstorm/nirstorm).

### 2.4. Standalone version

An executable, platform-independent software based on Java is available for download on Brainstorm website and enables to perform fNIRS analyses without requiring a MATLAB® license. The only feature that is not available in this compiled version is the optimal montage module (section 5), which relies on CPLEX Optimization software^32^.

## 3. fNIRS optimal montage design and personalized fNIRS investigation

### 3.1. The concept behind the fNIRS optimal montage

A critical step when performing an fNIRS experiment is the choice of optode positions on the scalp, especially when using fNIRS devices allowing only partial spatial coverage. In fNIRS, a standard sensor configuration consists of placing the optodes using a grid with a fixed distance of about 3 cm between the sources and the detectors over a specific region of interest (ROI). However, since light propagation in brain tissues is non-trivial, using only such a simple geometric criterion might not lead to a montage with the optimal sensitivity to the targeted cortical ROI. Additionally, it has been shown that a high-density montage could improve the reconstruction of hemodynamic fluctuations on the cortical surface^33^. However, because the number of sources and detectors available is often limited, a tradeoff must be found between featuring a local high-density montage and a sparse montage covering a more extended spatial region.

To address this issue, we proposed an original “optimal montage” methodology, which determines the position of a set of optodes that maximizes montage sensitivity over a targeted ROI while accounting for possible geometrical constraints imposed by the user on the position of the optodes^14,34^. The originality of the method consists in specifying a constraint on the ‘adjacency number’, which is the minimum number of channels created for each light source. The objective of this constraint is to allow the creation of spatially overlapping channels, to improve the accuracy of subsequent 3D reconstruction methods^31,35^. Other specific constraints may be added, such as the need to keep specific positions available to allow the installation of EEG electrodes according to a standard EEG montage, for simultaneous EEG/fNIRS experiments^14^.

### 3.1. Mathematical formulation of the fNIRS optimal montage

The procedure to compute a personalized fNIRS optimal montage consists of the following workflow presented in Cai et al 2021^31^:

1. Defining the target region of interest (ROI) on the cortical surface.
2. Defining the search space containing the set of possible optode positions along the high-density tessellated surface of the scalp (∼ 4000 vertices, edge length ∼ 5 mm) segmented from the anatomical T1-MRI of the subject or from a template (Figure 1.B). By default, the search space contains all the vertices on the scalp within a specific distance to the targeted region (e.g., less than 5 cm apart). However, this region can be fine-tuned either to allow simultaneous EEG-fNIRS acquisition by removing specific locations dedicated for EEG electrode installation from the list of locations of fNIRS optodes or when considering an fNIRS cap by allowing only sparse possible locations of fNIRS sensors.
3. Performing Monte Carlo simulations for near-infrared photon transport into biological tissues using MCXLAB software^36^ to generate “all” possible light sensitivity profiles (∼200,000 sensitivity profiles generated in the example presented in Figure 1).
4. Solving a mixed linear integer problem given specific functional constraints. (Figure 1.B). The optimization problem is formulated as a mixed linear integer programming problem under functional constraints, which is solved in NIRSTORM using IBM ILOG CPLEX Optimization toolbox^37^ (IBM, version 13, https://www.ibm.com/products/ilog-cplex-optimization-studio). The constraints include constraints on the equipment available (number of sources, and detectors), distance constraints (minimum source-detector distance, and range of distance between any optodes), and adjacency constraints (minimum number of channels formed by each source).

**Figure 1.**
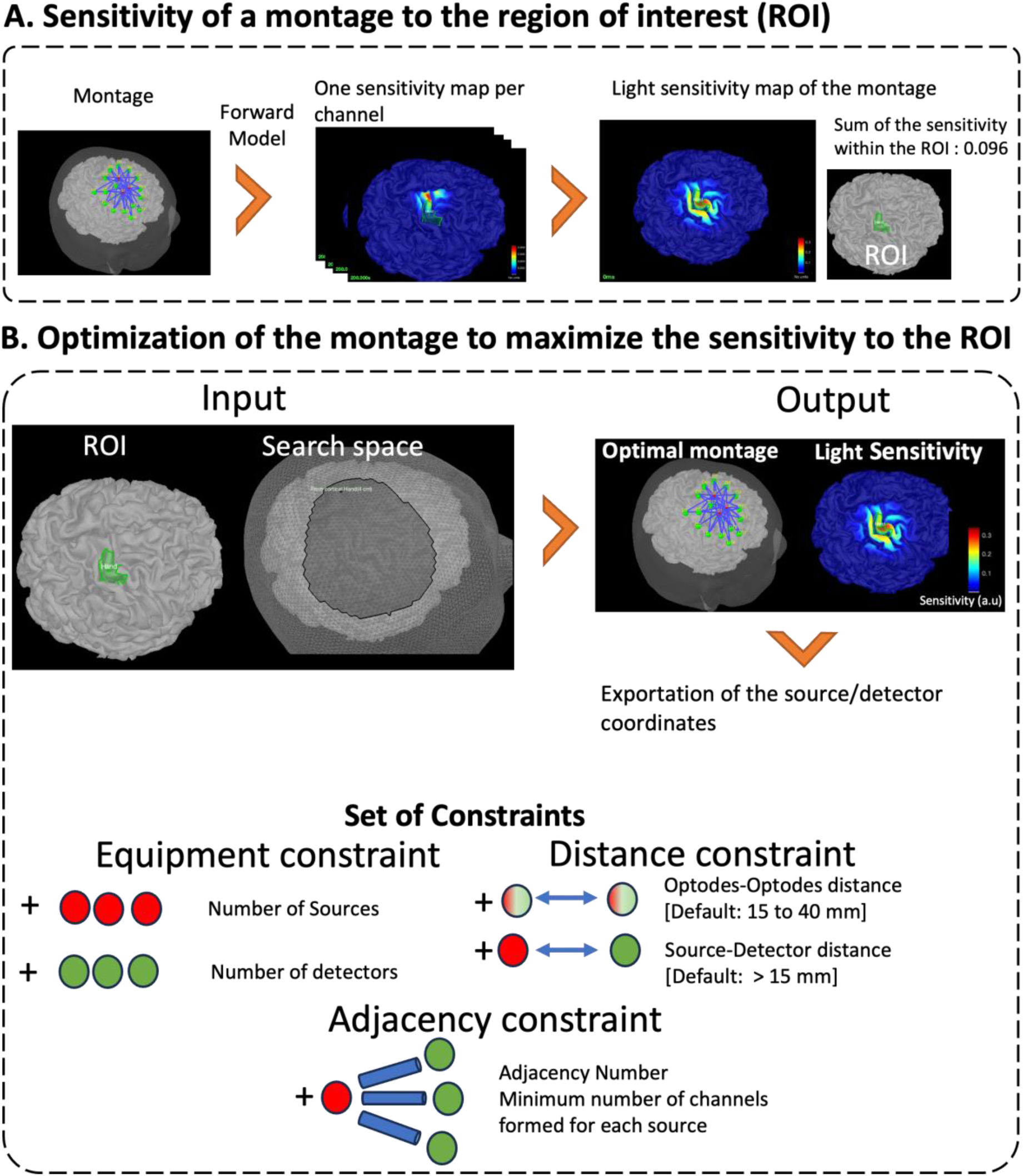
Description of the optimal montage. A. Determination of the light sensitivity of a specific montage to a region of interest (ROI) using the fNIRS forward model (sum of the light sensitivity within the ROI). B. Determination of the montage that maximizes that sensitivity based on the search space (all positions on the skin mesh) and a set of constraints on the montage.

Further details on the implementation can be found in Machado, et al. 2018^15^.

Finally, the optimal position of the optodes can be exported in different formats and coordinate systems to be used for data acquisition. The coordinates information can be used with a 3D neuronavigation system to guide the installation of the optodes on a participant’s head or to guide the fabrication of a specific fNIRS cap.

## 4. Standard channel space analysis of fNIRS signals

The overall objective of this section is to introduce standard fNIRS analysis conducted at the channel space including standard preprocessing, estimation of hemoglobin local variations using the modified Beer-Lambert law, and detection of brain activation using the general linear model.

### 4.1. Input data

The analysis of fNIRS data analysis requires the following input data at a minimum: (i) recorded fNIRS time series, (ii) channel configuration. More specific information can be added: (iii) anatomical data, (iv) description of the optical montage, i.e., the spatial location of the sources and detectors.

Functional data contains time series, information concerning the task and stimuli, and information regarding the fNIRS montage. NIRSTORM supports the two main data file formats used to store fNIRS data: Homer file format (.nirs)^38^ and the shared near-infrared file format specification (.snirf)^39^. To facilitate the coregistration of the optodes’ location on anatomy, NIRSTORM fully benefits from Brainstorm features for handling anatomical data^29,30^.

Anatomical information can be extracted from subject-specific MRI data (T1-or T2-weighted MR images) or can be provided by an anatomical template (e.g., Colin27^17^, ICBM152^18^). When using a subject’s specific MRI for head modeling, the fNIRS montage will be automatically coregistered to the subject’s anatomy using anatomical reference points (i.e., fiducial points). Additional head points digitized from the head surface of the subjects should be considered to improve the overall spatial coregistration of fNIRS sensor locations to the anatomical head model. A rigid transformation matrix (3 rotations, 3 translations) is estimated using the iterative closest point algorithm implemented in Brainstorm^29^ ensuring an accurate coregistration between the skin mesh segmented from the MRI and the digitized head shape.

For fNIRS forward models and visualization purposes, surface representations of the subject’s skin and cortical surface, as well as a volume segmentation of the whole head into five tissues are required (typically volume segmentation of skin, cerebrospinal fluid [CSF], skull, grey matter [GM], white matter [WM]). Preparation of these anatomical data can be obtained using well-known anatomical head modeling packages already integrated into Brainstorm (e.g., CAT12^40^, FreeSurfer^41^). When no individual MRI is available, an anatomical head model (segmented volumes and surfaces) from a template MRI can be scaled or deformed to fit the subject’s digitalized head shape.

### 4.2. Preprocessing

This section aims to describe the main preprocessing tools that should be considered for standard fNIRS analysis. The main preprocessing steps include bad channel detection, motion correction, frequency filtering, physiological noise regression, and computation of HbO and HbR fluctuations.

#### 4.2.1. Bad channel detection

Bad channel identification, sometimes called fNIRS channel pruning, aims to identify channels without reliable measurement that should be discarded from further analyses. This may correspond to channel measurements containing too noisy signals or channels contaminated with signals of non-interest (i.e., motion artifacts or equipment noise). Noisy channels could sometimes exhibit negative values (large variability around zero) and, therefore, could not be considered for modified Beer-Lambert conversion, which requires applying a logarithm. Our strategy is to discard those channels. Channels contaminated with signals of no interest can be detected by visual inspection or by using metrics such as the scalp coupling index^42^ (SCI) or the coefficient of variation^43^ (CV). SCI evaluates the quality of a channel by assessing the presence of cardiac oscillations in the signal. The underlying assumption is that a “good” channel should measure similar fluctuations in the cardiac band for each wavelength. SCI is obtained by bandpass filtering the data within the cardiac band (typically, 0.5 to 2.5 Hz) and then computing the correlation between the optical densities measured for each wavelength. CV measures the amount of signal variation in each channel by estimating the ratio between the standard deviation and the mean of the light intensity along time for each channel. Typical values of SCI less than 80% or CV larger than 30% would indicate noisy channels. Bad/noisy channels can then be excluded from further analyses. Both SCI and CV metrics are implemented in NIRSTORM.

#### 4.2.2. Motion correction

Motion is an important source of artifacts in fNIRS signals and must be handled/corrected carefully. Several approaches have been proposed in the literature to detect and correct motion artifacts^44^. Two standard approaches were implemented in NIRSTORM (Figure 2): (1) a semi-automatic algorithm that relies on a manual identification of motion artifacts followed by spline-interpolation^45^ (2) the Temporal Derivative Distribution Repair (TDDR) method^46^, an automatic motion correction algorithm that relies on the assumption that motion will introduce outliers in the temporal derivative of the optical density.

**Figure 2.**
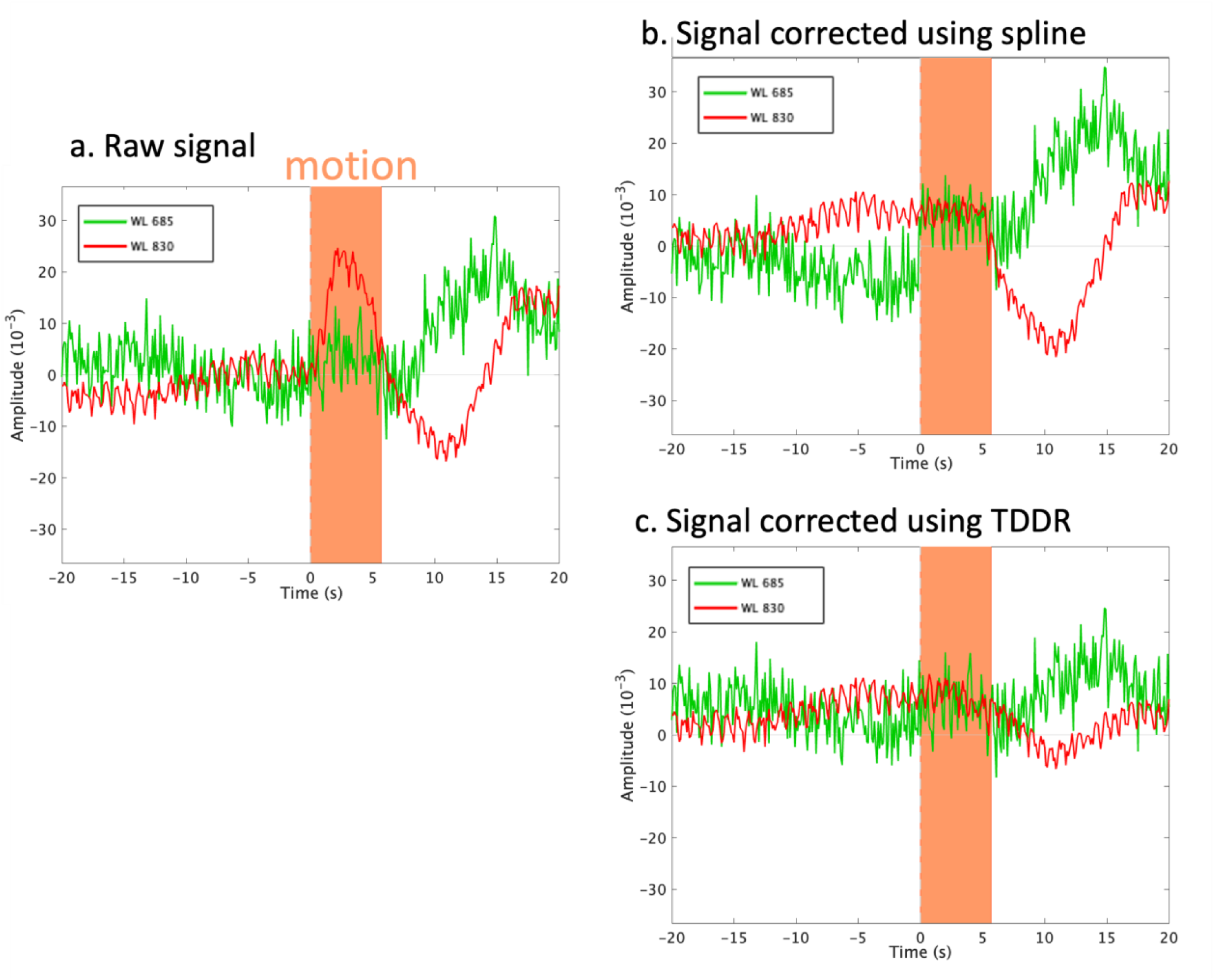
a. ΔOD signal for wavelengths 685nm (green) and 830nm (red) and manual identification of the motion window (in orange). b. fNIRS signal corrected using spline interpolation. c. fNIRS signal corrected using Temporal Derivative Repair (TDDR). The windows are centered around the detected motion (–20 s to 20 s). The identified motion lasted 5 seconds.

#### 4.2.3. Frequency filtering

Filtering can be used to improve signal quality by removing frequency bands of no interest, typically high frequencies containing physiological signals (e.g., Mayer waves, breathing, heartbeat) or instrumental noise, and by removing low frequencies corresponding to very slow signal drifts. Two options are available in NIRSTORM, using either a finite impulse response (FIR) or an infinite impulse response (IIR) filter (e.g., 3rd-order Butterworth filter). Additionally, NIRSTORM also provides a detrending function to remove slow fluctuations of non-neuronal origin from fNIRS signals. This usually consists of filtering out all frequency components featuring a period longer than 200 seconds (i.e., frequency components lower than 0.005 Hz) by regressing out a set of discrete cosines transform functions using a linear regression approach.

#### 4.2.4. Physiological noise regression using short separation channel regression

A typical source of physiological noise that can arise in fNIRS signals comes from the superficial layers of the head. Extracerebral physiological fluctuations of no interest are mainly driven by cardiac activity, breathing, blood pressure changes, and vasomotion^47^. However, because some of these very slow fluctuations overlap with the frequency bands of brain signals of interest, they cannot simply be removed using frequency filtering. Short separation channels (SSC) consist of adding fNIRS channels for which the source-detector distance would be around 0.8 cm in adults. SSC are designed to measure fNIRS signals from superficial layers only, i.e., not from the brain. Therefore, SSC signals can be used to regress out physiological fluctuations originating in superficial layers from channels of interest using a standard linear regression approach. It is worth mentioning that SSC will also be distorted by head motion, so this approach also allows additional motion correction of fNIRS signals. Several methods have been proposed for short-channel regression^48–51^. In default NIRSTORM implementation, a regression model is fitted for each chromophore/wavelength using an ordinary least squares approach (OLS), considering the mean of all superficial channels for that chromophore/wavelength to model the superficial signal.

#### 4.2.5. Estimation of HbO and HbR fluctuations using the modified Beer-Lambert law

Measured light intensities recorded using the fNIRS device should first be converted in fluctuations of optical density (**ΔOD**) as follows:

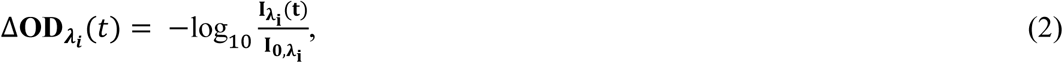

where, *I*_λ*i*_ represents the raw light intensity measured over time *t* for a specific channel for a particular wavelength λ_*i*_. *I*_0,λ*i*_ is a scaling factor used for baseline correction and computed for each channel using the mean (or median) value within a baseline window specified by the user.

Relative hemoglobin changes (**ΔHbO**, **ΔHbR**) can then be estimated for each channel using the modified Beer-Lambert law (MBLL) as follows:

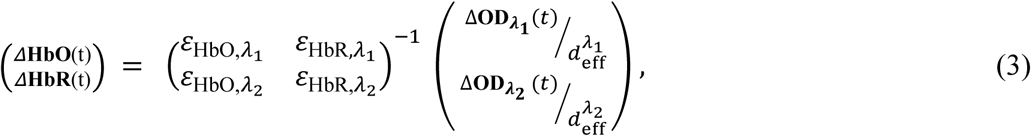

where ε_HbO,λ*i*_ and ε_HbR,λ*i*_ are respectively the extinction coefficient of the oxy– and deoxy-hemoglobin for a specific wavelength λ, and 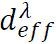 is the estimation of the effective light pathlength between the source and the detector at a specific wavelength λ:

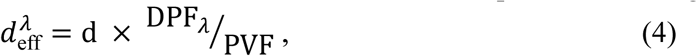

where d is the actual source-detector distance, DPF_λ_ is the differential path-length factor calculated based on the wavelength of interest and the age of the subject, using the formula described either in Duncan et al., 1996 ^52^ or Scholkmann et al, 2013^53^. DPF_λ_ typically ranges between 5 and 6. Finally, PVF is the partial volume factor, as defined in Scholkmann et al, 2013 ^53^, the default value being set to PVF = 50. 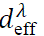 would typically range between 0.3 and 0.4 cm for a source-detector distance of 3 cm. Finally, once **ΔHbO** and **ΔHbR** are estimated, the total hemoglobin ΔHbT is then computed as the sum of **ΔHbO** and **ΔHbR**.

### 4.3. Standard hemodynamic response estimation using fNIRS signal averaging

A typical fNIRS task design would consist of repeating stimuli, following either a block or an event-related design, to increase the signal-to-noise ratio of the specific brain hemodynamic response elicited by the experimental conditions. Similarly to EEG event-related potentials, one can extract, for each fNIRS channel, the evoked hemodynamic response to a specific condition (block or event-related) by extracting and averaging epochs of data around the onset of each condition. Several standard tools developed for EEG event-related potentials analysis within Brainstorm can then be used to compute, visualize, and analyze these evoked hemodynamic fNIRS responses (Figure 3). As expected for a finger-taping task, the average hemodynamic response shows an increase of HbO and a decrease of HbR in the contralateral somatomotor region. Moreover, this averaged response can also be normalized (e.g., z-score normalization with respect to a baseline) and statistics can be computed to compare the magnitude of this response between subjects and conditions (see Tadel et al., 2019^29^ for a complete review).

**Figure 3.**
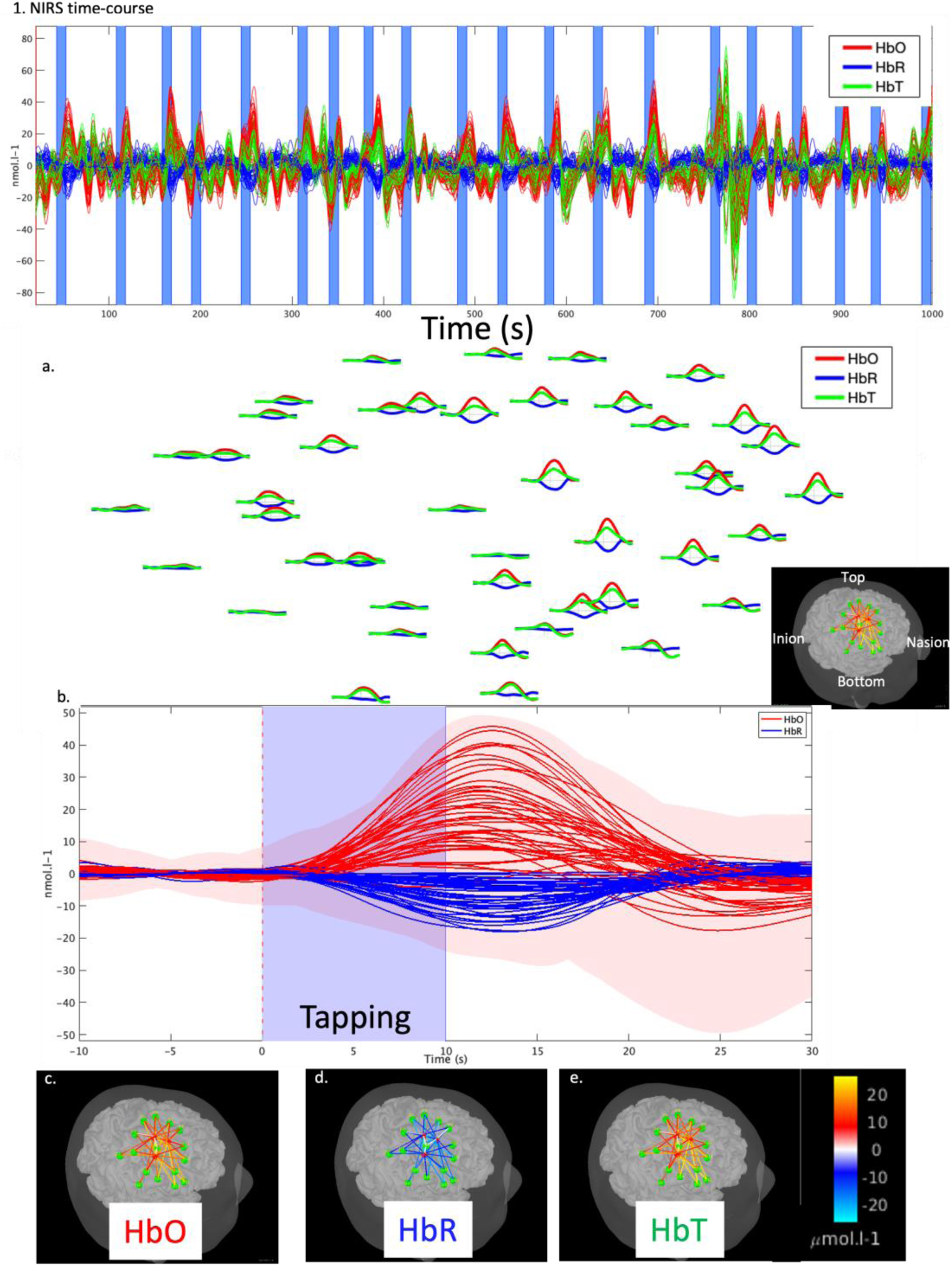
Standard channel-based analysis of fNIRS finger-tapping data using epoch averaging. (1) Hemodynamic time course for HbO (red), HbR (blue), and HbT (green); each channel is displayed as a solid line. Vertical blue shaded areas denote the finger-tapping periods. (2) Averaged evoked hemodynamic response using three different visualizations of the evoked hemodynamic response: a. 2D layout: evoked hemodynamic response for each channel. As expected for a right-finger-taping task, the average shows an increase in HbO and a decrease in HbR in the somatomotor region contralateral to the finger used. b. evoked hemodynamic response time course for all channels. c,d,e. Topographies of the hemodynamic response at t=+11.9 s post-stimulus for HbO (c), HbR (d), and HbT (e), where signal amplitude is represented along each channel (represented by a cylinder)

### 4.4. Detection of brain activation using the general linear model

Since fNIRS measures hemodynamic signals, the expected hemodynamic response features similar properties to fMRI responses elicited by a task. Therefore, we also implemented a statistical module allowing users to analyze data using the classical general linear model (GLM), reminiscent of fMRI analysis (see Monti, 2011^54^ for a review) to detect brain activation. The general linear model is specified at the subject’s level as follows:

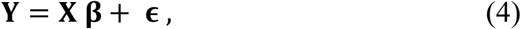

where **Y** is the measured data for a specific channel (for HbO, HbR, and HbT separately), **X** is the design matrix containing the regressors of interest and potential confounds. β contain the estimated weights associated with each regressor and ϵ is an error term. Within the design matrix, the expected hemodynamic response of interest is computed by convolving the time course of the task paradigm (block, event-related) with a predefined canonical hemodynamic response function (HRF). The canonical HRF model proposed in NIRSTORM is shown in Figure 4.3 exhibiting a peak at 5 seconds, and an undershoot at 15 seconds, as proposed by Glover et al^55^. It is important to note that the same HRF is used to model the hemodynamic response for HbO, HbR, and HbT, which could be considered as a limitation since HbO is known to peak earlier than HbR in physiological conditions^56^. Implementing more advanced HRF models dedicated to fNIRS, such as spline basis function^57^ will be considered in future releases. Within the design matrix, we could add additional nuisance regressors to model hemodynamic fluctuations unrelated to the paradigm. In this context, NIRSTORM allows the user to include as additional confounds discrete cosine-basis functions (see section 3.1.3) to regress out slow drifts in the signals as well as SSC data to regress out physiological fluctuations from superficial layers. One difference with other software packages is that the regressor used for short channels differs between HbO, HbR, and HbT: the HbO signal of short channels is used as a regressor only for the HbO signal.

**Figure 4.**
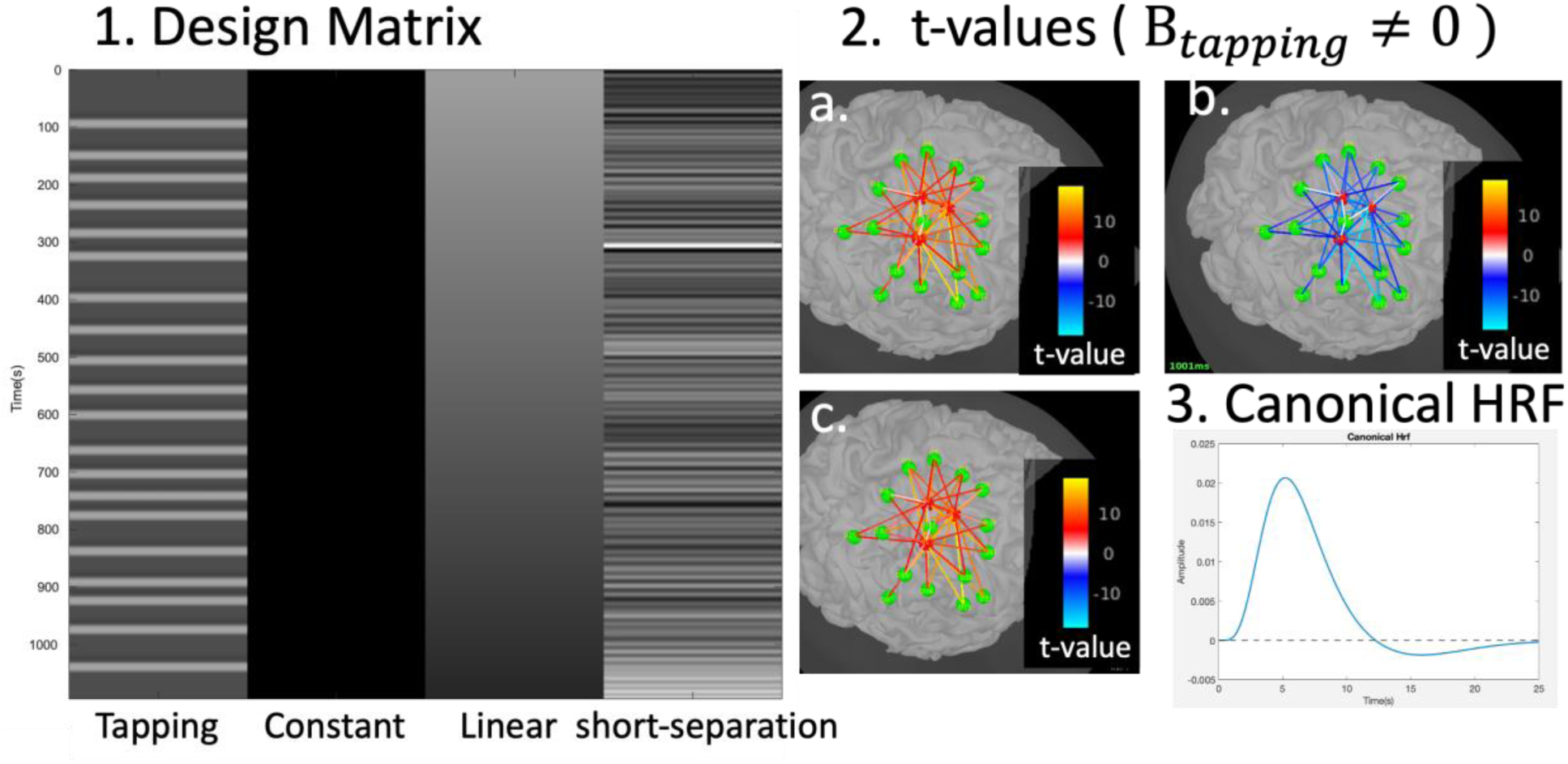
General Linear Model. 1. Design matrix of the model: the proposed model contains 4 regressors, from left to right: the task regressor, convolved with a canonical HRF, a constant, a linear trend, and the mean of SSC signals. 2. Result of the Student t-test for the contrast +β_*tapping*_ ≠ **0** for HbO (a), HbR (b) and HbT (c). The color scale shows the Student’s t values. In this example, all the channels cover the active region and are all statistically significant (p < 0.05, FDR corrected). 3. Canonical HRF used in NIRSTORM.

The error term ϵ is usually assumed to be Gaussian and independently distributed. However, temporal autocorrelations are present in the time series. In NIRSTORM, we implemented two strategies to model these temporal autocorrelations between fNIRS samples: the precoloring and the prewhitening approaches. The linear model can then be rewritten as ***SY*** = ***SX*** β + ***S***ϵ, where **S** can be obtained either by precoloring the data and the model (where **S** is a filter based on the canonical hemodynamic response) or by prewhitening the data and the model (where ϵ is modeled using an autoregressive model of order 1, the corresponding noise covariance structure being then considered for the prewhitening). NIRSTORM implementation follows the models proposed by Tak et al, 2014^58^

Once the serial autocorrelation is modeled using either the precoloring or prewhitening strategies, it is possible to estimate the parameters β, for each channel, using ordinary least-squares fitting, as follows:

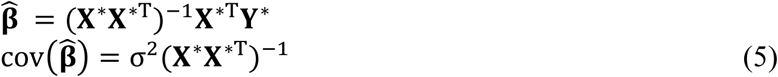

where ***X***^∗^ = ***SX***, ***Y***^∗^ = ***SY***, σ^2^ is the variance of the residual (***S***ɛ), and ^T^ denotes matrix transposition.

Once parameters β are estimated for each channel, statistical contrasts (i.e., linear combinations of parameters) can be defined to compare the brain activations elicited by different experimental conditions.

Statistical significance can then typically be assessed using the Student t-test on the defined contrasts. The statistical test can be performed either at the subject or group levels. The group-level effect is estimated using a mixed-effect model as described in Tak & Ye, 2014 ^58^. Finally, a p-value threshold can then be applied to the resulting statistical maps, which can be corrected for multiple testing using either Bonferroni or false discovery rate (FDR) approaches^59^.

It is worth mentioning that in NIRSTORM the GLM module can be used either in the channel space or after 3D reconstruction along the cortical surface.

Figure 4.1 shows the design matrix X containing 4 regressors: the task regressor, convolved with the canonical HRF (figure 4.3), a constant, a linear trend, and the mean of SSC signals. Figure 4.2 shows the result of the two-sided Student t-test for the task-related contrast. In this example, all the channels covered the active region, consequently, for these high SNR data, all channels exhibited statistically significant responses (p < 0.05, FDR corrected) and showed, as expected, an increase in HbO and HbT (positive t-value) and a decrease in HbR (negative t-value).

## 5. fNIRS 3D reconstruction using Near Infrared Optical Tomography

Near-infrared optical tomography (NIROT) consists in using fNIRS measures of optical density from at least two wavelengths to infer 3D reconstructions of local fluctuations of HbO and HbR along the cortical surface. Solving NIROT 3D reconstruction amounts to solving an ill-posed inverse problem^60,61^, which is very similar to the EEG/MEG source imaging inverse problem^62^.

### 5.1. Forward model of NIR light propagation within the head tissues

Before solving the NIROT inverse problem, one needs to solve the forward model describing the propagation of NIR light within head tissues in a realistic manner. The forward model consists of estimating how local fluctuations of NIR light for each wavelength λ on the cortex (which are due to differential light absorption HbO and HbR) affect scalp-level fNIRS measurements. To solve this problem, we need a model describing how light propagates inside the head from each fNIRS source to each fNIRS detector. The fluences from a specific source to a specific detector are estimated using realistic Monte Carlo simulations of light photon transport, using the photons transport simulator implemented in MCXLab toolbox^36^. To facilitate its usage, we integrated MCXLab toolbox as a plugin inside Brainstorm environment. For each channel, the propagation of millions of photons through the head should be simulated (usually between 10 to 100 million photons). The head is usually modeled using a 5-tissues layer description (skin, skull, cerebrospinal fluid, grey matter, white matter) obtained from the segmentation of anatomical MRI data^41^, which allows to set the optical properties of each tissue. Once all fluences are computed, for all source-detector pairs, a volumetric light sensitivity map can be estimated for each channel, which assesses how a local change in HbO/HbR would impact fNIRS measurement at the channel level. Volumetric light sensitivity maps are computed by using the adjoint method according to Rytov approximation^61^. It is worth mentioning that fluence estimation is computationally intensive, especially when several channels are considered along a detailed cortical surface. Therefore, GPU implementation is recommended, but not mandatory (see http://mcx.space/wiki/?Speed for a comparison between CPU vs GPU-based implementations).

The forward model **A**_λ_is a matrix (number of channels x number of vertices) estimating the contribution of a local change in optical density (caused by changes in HbO/HbR concentration at a specific vertex on the cortical surface) to the observed change on a specific channel on the scalp, for wavelength λ. **A**_λ_ is obtained by projecting the volumetric sensitivity map obtained using MCXLab to the cortical surface. Such projection is achieved using a Voronoi projection: each voxel is associated with a Voronoi cell describing the interpolation kernel for a specific vertex of the cortical surface, resulting in an anatomically informed interpolation scheme based on Voronoi diagrams^63^. The resulting volume-to-surface interpolation scheme is therefore informed by the underlying anatomical sulco-gyral geometry. The projection is completed by estimating the sensitivity of each vertex as the mean light sensitivity within each Voronoi cell.

### 5.2. Reviewing the fNIRS forward model

Importantly, the resulting forward model, mandatory for NIROT, can also be used to assess how the montage is spatially sensitive to specific cortical regions (Figure 5). Such an approach is also important when considering a sparse fNIRS montage, not designed for 3D reconstructions, but allowing an accurate assessment of which cortical brain regions could be monitored with a specific montage. A light sensitivity map can be represented for each channel, or for the entire montage by summing the sensitivity maps of all channels. Because the unit of the sensitivity matrix is not directly interpretable, sensitivity maps in NIRSTORM are converted to decibels (dB), so that 0 dB corresponds to the vertex of the cortical surface exhibiting maximal sensitivity. Therefore, –1 dB corresponds to a vertex for which the fNIRS montage is 10 times less sensitive when compared to the vertex exhibiting maximal sensitivity. Since spatial overlap between fNIRS channels is important before considering NIROT reconstruction, we also added two measures of spatial overlap in NIRSTORM. To estimate the spatial overlap between fNIRS channels, a threshold must first be set, considering either the minimum number of channels sensitive to a specific cortical region or the minimum sensitivity required to receive signals from that region.

**Figure 5.**
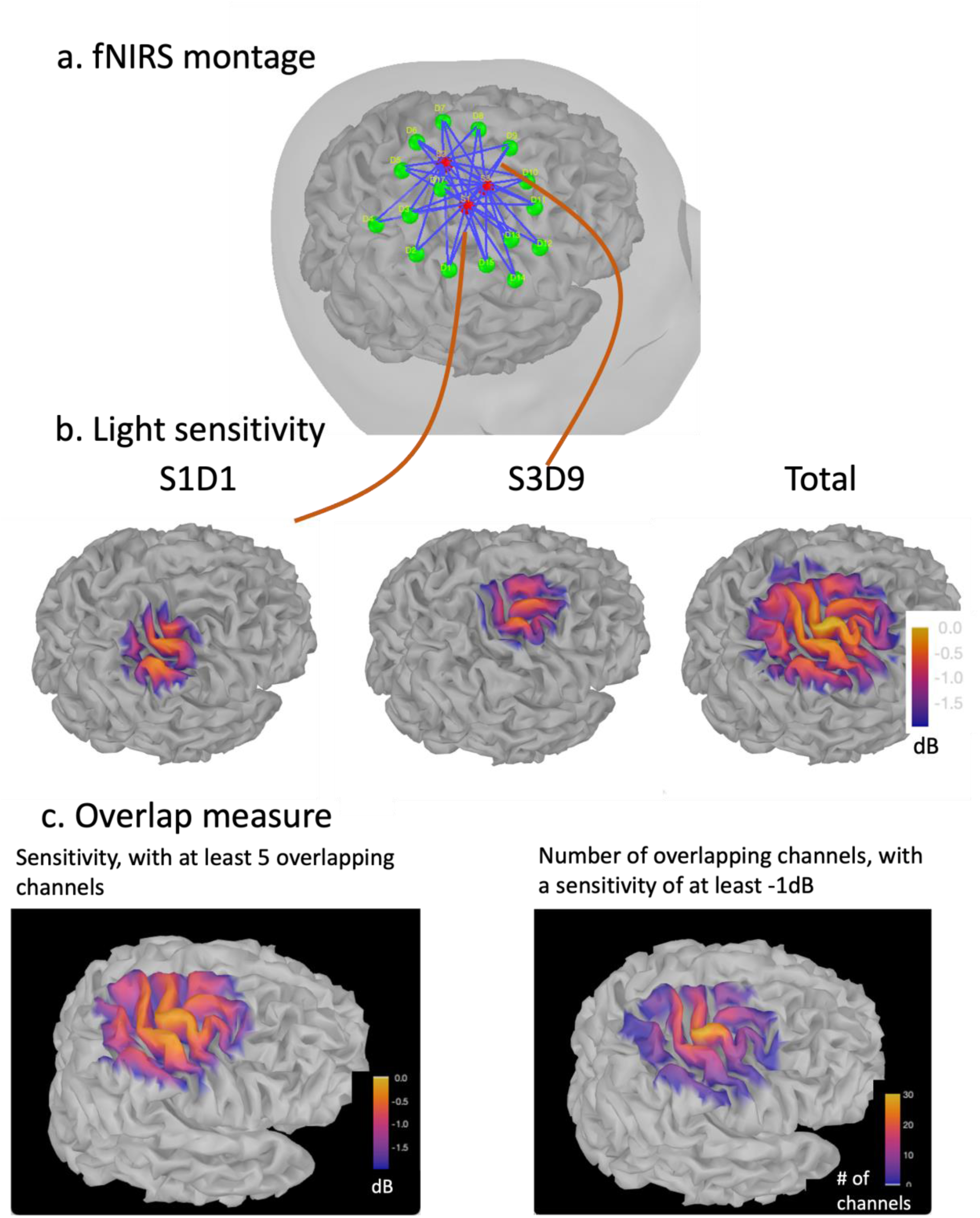
fNIRS forward model estimation. A. Illustration of the fNIRS optimal montage. (sources in red, detectors in green, and channels in blue). b. Light sensitivity maps of two channels (S1D1 and S3D9) covering different regions, as well as the light sensitivity map of the whole montage. Yellow indicates regions exhibiting higher sensitivity. C. Spatial overlap measure expressed as sensitivity involving at least 5 overlapping channels, or the total number of spatially overlapping channels exhibiting at least a –1 dB sensitivity.

When setting the threshold to the expected minimum number N_*c*_ of channels sensitive to a region, NIRSTORM reports, for each vertex, the minimum sensitivity achieved by the N_*c*_ most sensitive channels. On the other hand, when specifying a threshold on the expected minimum light sensitivity, the algorithm reports for each vertex how many channels are sensitive to this specific vertex (above the specified threshold). We believe this is an essential tool for investigating the spatial sensitivity profile of fNIRS montage (sparse versus dense montages) while allowing us to assess whether sufficient spatial overlap between channels is reached before considering NIROT 3D reconstruction.

### 5.3. Solving NIROT Inverse problem

The inverse problem aims at reconstructing the fluctuations of fNIRS hemodynamic signals along the cortical surface (HbO, HbR) based on the optical density measured on the scalp at the channel level. NIROT involves two steps: first, optical density fluctuations are estimated along the cortical surface from scalp recordings (sections 5.3.1 and 5.3.2). Second, hemodynamic fluctuations in HbO and HbR are estimated on the cortical surface from the reconstructed local changes in optical density at two wavelengths, using Beer-Lambert law (section 5.3.3).

Solving NIROT problem aims at solving the following linear problem, which needs to be inverted for each wavelength separately:

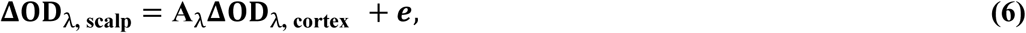

where **ΔOD_λ, scalp_** is an N_*c*_ × N_*t*_ matrix of scalp measurements recorded on N_*c*_ channels (i.e. N_*c*_ source-detector pairs) at N_*t*_ time samples for a specific wavelength λ. **A**_λ_ is the N_*c*_ × N_*v*_ matrix containing the forward model. In practice, only the vertices located within the field of view of the fNIRS montage should be considered for NIROT. This can be done by selecting all the vertices located within less than for instance 3 cm from any source or detector. **ΔOD_λ, cortex_** is an unknown N_*v*_ × N_*t*_ matrix containing the fluctuations over time of the optical density at wavelength λ_*i*_ along N_*v*_ vertices of the cortical surface. ***e*** is a N_*C*_ × N_*t*_ noise matrix corresponding to the additive measurement noise.

**Notations:** Since the inverse problem is solved independently for each wavelength, and to simplify the equations, we will consider the following simplified notations: ***m***(*t*) will denote the data recorded on the scalp **ΔOD_λ, scalp_** (*t*), the fluctuations of optical density along the cortical surface **ΔOD_λ, cortex_** (*t*) will be denoted ***j***(*t*), and the forward model **A**_λ_ will be denoted ***A***.

In NIRSTORM, we implemented and validated two NIROT methods: (1) Minimum Norm Estimate (MNE)^64–66^ and (2) coherent Maximum Entropy on the Mean (cMEM)^31,35,67–69^.

#### 5.3.1. Minimum Norm Estimate

The Minimum Norm Estimate (MNE) is a linear method providing a distributed solution that exhibits minimum energy. Such an approach is also well known as the Tikhonov regularization approach ^10,64^.

MNE solution can be obtained by solving the following optimization problem:

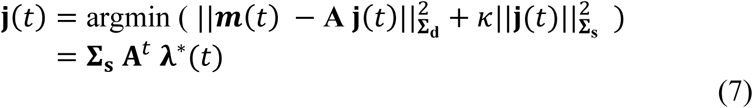

With λ^∗^(*t*) a linear function of the data ***m***(*t*) of dimension N_*c*_ given by:

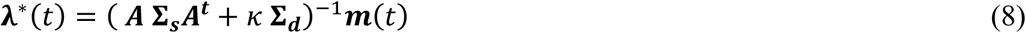

where κ is a regularization hyper-parameter, which can be estimated using the L-curve approach^70^. **Σ_d_** (N_*c*_ × N_*c*_) and **Σ_s_** (N_*v*_ × N_*v*_) are the covariances matrix at the channel level and along the cortical surface. **Σ_d_** is estimated as a diagonal matrix characterizing the noise covariance estimated from a baseline period. For standard MNE, the prior covariance **Σ_s_** of the fluctuations of optical densities along the cortex is usually chosen as the identity matrix. However, all uncertainties are not homogenous on the cortex, since signals generated in deeper regions will exhibit greater uncertainty. Therefore, an a priori source covariance matrix for the fluctuations of optical densities along the cortex should appropriately account for the variance differences across different brain locations and depths. To do so, a depth-weighted extension of MNE solution has been implemented in NIRSTORM. In this context, **Σ_s_**, the covariance matrix for the fluctuations of optical densities along the cortex, is defined by the following diagonal coefficients: 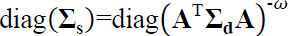 where ω is depth-weighted factor ranging from 0 (no depth weighting) to 1. After evaluating MNE NIROT accuracy using realistic simulations, Cai et al^35^ suggested that setting ω to between 0.3 and 0.5 would result in optimal fNIRS reconstructions using depth-weighted MNE.

#### 5.3.2. coherent Maximum Entropy on the Mean (cMEM)

The coherent Maximum Entropy on the Mean (cMEM) method is part of the Brain Entropy in Space and Time (BEst) toolbox, implemented as a plugin for Brainstorm software. The Maximum Entropy on the Mean (MEM) framework was first proposed by Amblard et al (2004)^67^ and then applied and evaluated by our group in the context of EEG/MEG source imaging^69,71^. MEM framework was specifically designed and evaluated for its ability to localize the underlying generators of EEG/MEG data along the cortical surface while featuring the unique property of recovering accurately the spatial extent of the underlying generators^72–77^. cMEM offers an efficient nonlinear probabilistic Bayesian framework to incorporate prior knowledge in the solution of the inverse problem. A key modeling feature is to consider a spatial prior model (or reference distribution), assuming that brain activity is organized within a set of *K* non-overlapping, and independent parcels, where the activity of each parcel is controlled by a hidden state variable. These non-overlapping parcels ^68^ are estimated using a data-driven parcellation of the full field of view considered for NIROT. While fitting the data through relative entropy maximization, cMEM has the unique ability to switch off parcels of the model considered inactive using a latent variable.

In our previous studies in the context of EEG/MEG source imaging, we have demonstrated the ability of cMEM to be sensitive to the spatial extent of the underlying generators, when dealing with focal sources^78^, as well as spatially extended generators in the context of clinical epilepsy data^76,77,79–81^. Moreover, we also demonstrated the excellent accuracy of cMEM in low signal-to-noise ratio (SNR) conditions, with the ability to limit the influence of distant spurious sources^77,78,82^. Accuracy in recovering the underlying spatial extent and robustness in low SNR conditions are two important properties to consider for fNIRS 3D reconstruction, especially in the context of maintaining a good intra/within-subject consistency using fNIRS due to its relatively low SNR^83^. We recently adapted and validated cMEM for NIROT reconstruction^31^, before applying such methodology to assess fNIRS response to changes in brain excitability elicited by transcranial magnetic stimulation^84^. Methodological details on cMEM formulation to solve the NIROT problem are described in Appendix A, whereas further details and validation are reported in our previous studies^31,81^.

Importantly, in Appendix A we are showing that the cMEM solution can be written, for each parcel

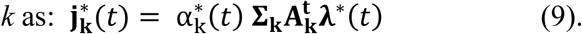

This equation is very similar to the equation of MNE (eq 7) with two important differences:

1. 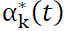 corresponds to the final probability of activation of each of the parcel *k* modulating the activity of each vertex within the parcel. **Σ**_**k**_ is the optical densities covariance of parcel *k*.
2. Unlike in MNE, there is no analytical expression for the N_*c*_ dimensional vector λ^∗^(t). Instead λ^∗^(t) is a nonlinear function depending on our prior and the data.

#### 5.3.3. Computing HbO and HbR fluctuations along the cortical surface

Finally, once local changes in optical density at each wavelength are reconstructed along the cortical surface, local hemodynamic fluctuations can be recovered along the cortical surface using the Beer-Lambert law (equation 10).

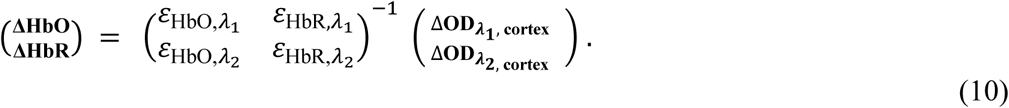

Since the inverse problem has already been solved and since **ΔOD_λ_**, **_cortex_** is now local, we do not need to consider the differential path length correction anymore.

A visual qualitative comparison between MNE and cMEM can be found in Figure 6 showing the averaged optical density response elicited by the finger-tapping task (Figure 6.1), the averaged hemodynamic response reconstructed within the hand-knob region for MNE and cMEM (Figure 6.2) and the estimated maps of HbO and HbR fluctuations at the time of the peak using MNE and cMEM. It can be noted that: (i) the estimated map obtained with cMEM is more focal and contrasted than that obtained with MNE showing a better estimation of the spatial extent of the underlying generators, (ii) MNE and MEM can both recover the hemodynamic response inside the motor region, both exhibiting ∼1s delay between HbO and HbR hemodynamic peaks

**Figure 6.**
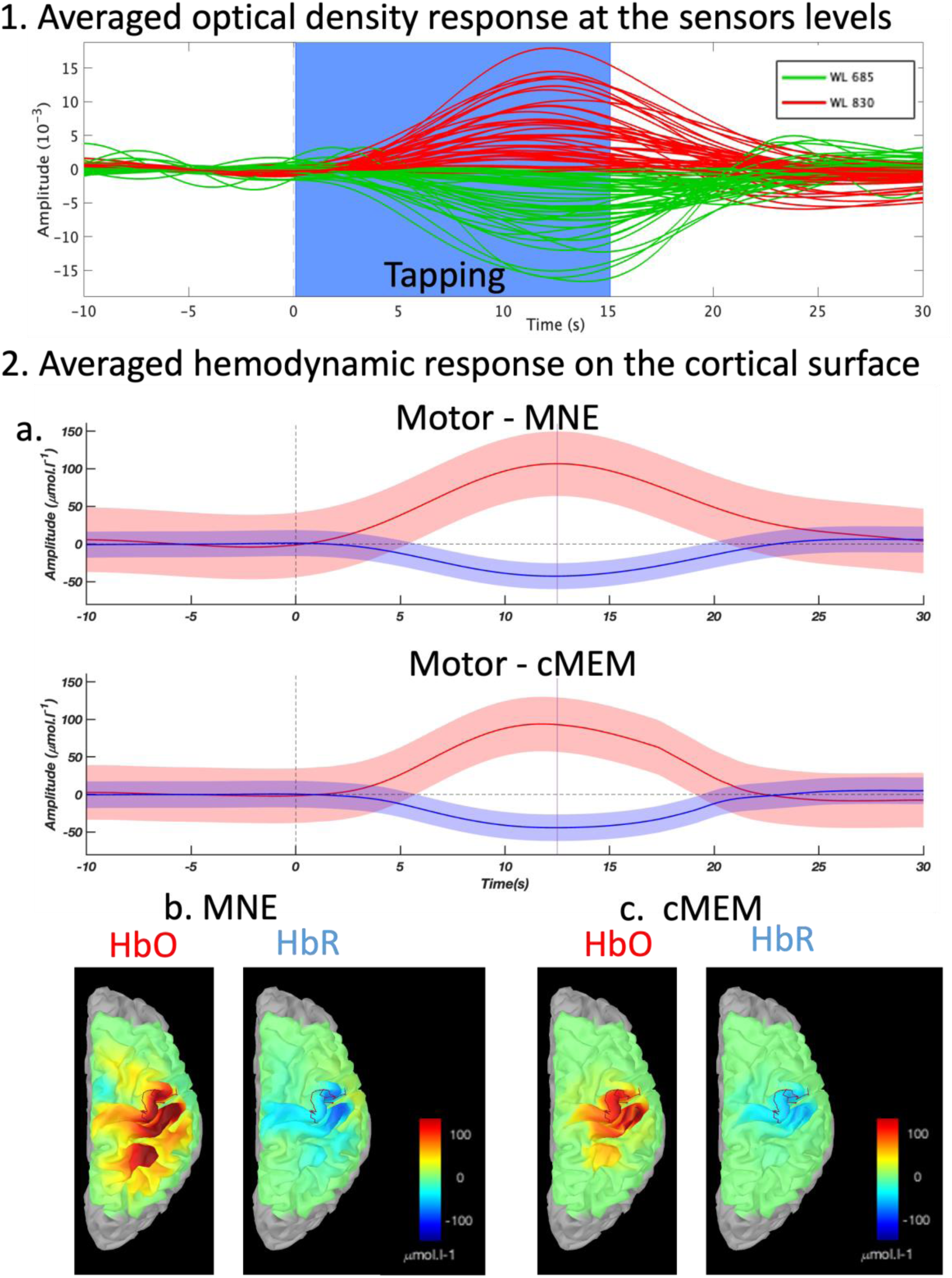
NIROT reconstruction module. 1. Averaged optical density response elicited by the finger-tapping task. The blue rectangle represents task duration. Each line corresponds to a channel (red corresponds to the 830-nm wavelength; green corresponds to the 685-nm wavelength). 2. Averaged hemodynamic response reconstructed within the hand-knob region using MNE and cMEM (and standard deviation across vertices) and estimated map at the time of peak for HbO and HbR using (b) MNE and (c) cMEM.

## 6. Multimodal integration – illustration in the clinical context of epilepsy

In this last section, we provide a typical illustration of how to leverage several tools and functionalities provided in the NIRSTORM/Brainstorm environment to conduct a complete multimodal analysis within the same unified environment. To do so, we present a study intending to investigate, using fNIRS, the interactions between sleep and epilepsy, within the presumed epileptic focus that has been localized using EEG/MEG investigation. In this study, we considered multimodal data collected on a patient with left frontal lobe epilepsy who underwent simultaneous EEG-MEG recording to localize the presumed epileptic focus. Personalized EEG/fNIRS investigation targeting the estimated epileptic focus and its corresponding homologous contralateral region was then considered for whole-night monitoring, following the method we proposed in Pellegrino et al 2016^7^.

These studies were approved by the Research Ethics Board of the Montreal Neurological Institute and Hospital for the EEG-MEG and by the Research Ethics Board of PERFORM Center (CCER 18-19 – 02) for the EEG/fNIRS whole night monitoring. A written informed consent was signed by the participant.

First, EEG-MEG data were analyzed within the Brainstorm environment^79^ to localize and delineate the patient-specific epileptic focus on the cortical surface, following the methodology described in Heers et al, 2014^73^. MEG interictal epileptic discharges (IEDs) were visually identified and marked at their peak by a trained epileptologist.

Corresponding EEG/MEG signals were then segmented into 2-second-long epochs around the interictal epileptic discharge peaks (−1 to +1s) and then averaged and localized on the cortical surface using cMEM, resulting in an EEG source imaging map and a MEG source imaging map (Figure 7.1.a). The activated regions detected using EEG and MEG source imaging were used to delineate the suspected epileptic focus (Figure 7.1.c). This information was then used to define the targeted regions to estimate an optimal montage for personalized fNIRS investigation, namely: the suspected focus, the corresponding contralateral homologous regions (later called the homologous region), and a control region (Figure 7.2.a) as proposed in Pellegrino et al 2016^7^.

**Figure 7.**
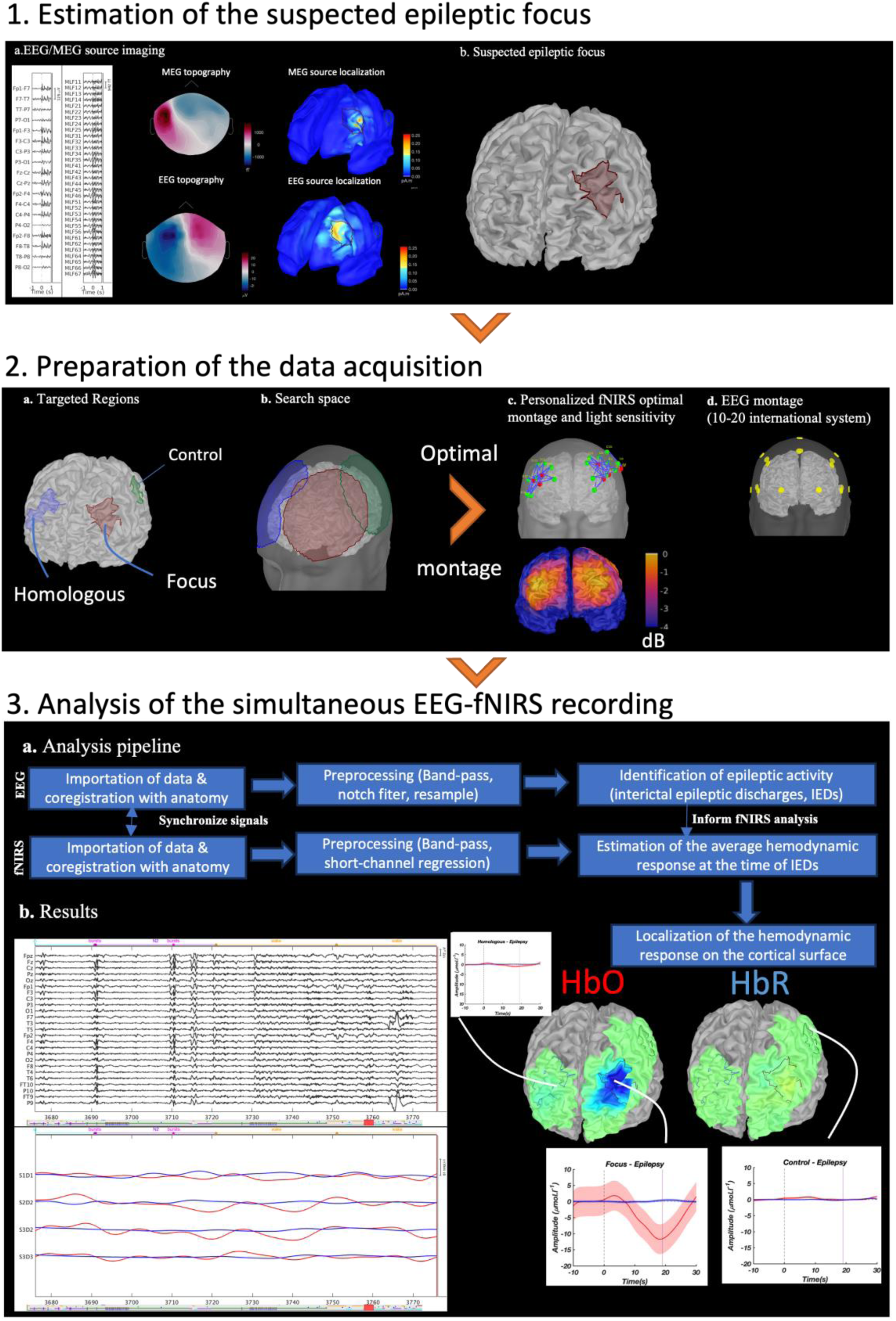
Multimodal investigation of a patient with left-frontal focal drug-resistant epilepsy. 1. Estimation of the suspected epileptic focus. 1.a. Simultaneous EEG/MEG recording showing a bilateral burst of polyspike, and waves and corresponding EEG/MEG source imaging results 1.b. Suspected left frontal epileptic focus based on MEG source imaging results. 2. Estimation of the fNIRS optimal montage targeting the suspected epileptic focus, the corresponding homologous contralateral region, and a control region not involved in the epilepsy. 2.a. Definition of the targeted regions. 2.b. Definition of the possible locations of the optodes for each targeted region. 2.c. Personalized fNIRS optimal montage and corresponding light sensitivity maps. 2.d. EEG montage used to record epileptic activity based on the international 10-20 system. 3. Simultaneous analysis of the EEG and fNIRS data during interictal epileptic spikes. 3.a. Summary of the pipeline used for simultaneous analysis of fNIRS and EEG data. 3.b. Simultaneous visualization of the EEG and fNIRS signals showing several bursts of polyspike and waves in EEG along with the corresponding synchronized fNIRS fluctuations. Hemodynamic response in the targeted regions and the estimated cMEM map at the time of the peak for HbO and HbR occurring 19 s after the interictal spikes detected on scalp EEG (spatial mean and standard deviation across the vertices in the ROI).

The search space for the possible positions of the optodes was therefore defined as all the vertices of the skin mesh within a 5 cm radius of the targeted regions on the cortex (Figure 7.2.b). We then estimated an optimal fNIRS montage for each of the three targeted regions using the following configuration: 3 sources and 7 detectors for the focus and homologous regions, and 2 sources and 2 detectors for the control region. The three montages were then merged, and the associated light sensitivity map was computed using the fNIRS forward model previously described (Figure 7.2.c). Additionally, the position of the optodes was compared to the position of the EEG electrodes based on the international 10-20 system to ensure that there would be no conflict between the EEG and fNIRS sensors installation. The optimal positions of the EEG and fNIRS sensors were then marked directly on the head of the subject using the Brainsight neuronavigation system, after coregistration of the patient’s MRI data to the patient’s physical head. fNIRS optodes were glued to their optimal position on the head of the subject using a clinical adhesive called collodion. The actual positions of the fNIRS optodes and EEG electrodes were digitized using the same neuronavigation system considered for the coregistration with the subject’s anatomical data. The positions of 100 randomly selected head points were also digitized to improve coregistration accuracy. EEG/fNIRS data were recorded simultaneously during a whole night (10 hours) to investigate the interactions between sleep and epilepsy, using an fNIRS personalized montage informed by EEG/MEG localization of the presumed epileptic focus. EEG data were recorded with a BrainAmp amplifier (Brain Products, Munich, Germany) with a sampling rate of 1 kHz. fNIRS data were recorded with a Brainsight fNIRS system (Rogue Research Inc, Montreal, Canada) using 8 sources, 16 detectors, and 2 proximity detectors at a sampling rate of 10 Hz. During data acquisition, Eprime software^85^ was used to send simultaneous triggers to both EEG and fNIRS systems, to allow accurate temporal synchronization.

EEG and fNIRS data were imported into Brainstorm. EEG and fNIRS sensors were first aligned with the subject’s anatomy by identifying three anatomical fiducial points in the patient’s anatomical MRI (nasion, left pre-auricular, right pre-auricular). Coregistration with the anatomical MRI was further refined using the additional headpoints. fNIRS and EEG data were synchronized in time using the Eprime triggers^85^. EEG was preprocessed inside Brainstorm and analyzed by a neurologist to identify transient spontaneous interictal epileptic discharges (discharges lasting few hundred milliseconds with a high SNR). fNIRS data were preprocessed using NIRSTORM, including conversion to optical density, bandpass filtering between 0.002 Hz and 0.1 Hz, and regression of the superficial noise using short-separation channels. Data were also visually reviewed to mark any potential contamination from motion artifacts. Then, the hemodynamic response to interictal epileptic discharges was estimated by extracting **ΔOD_λ*i*, scalp_** signal around each isolated interictal epileptic discharge [-10 s to 30 s] (an interictal epileptic discharge was classified as isolated if no other epileptic discharge occurred in the window from –10 s to 30 s around the peak of the discharge). Epileptic discharges occurring at the same time as motion artifacts were discarded from further analysis. The corresponding channel level fNIRS averaged hemodynamic responses were finally localized along the cortical surface using cMEM. The estimating fNIRS hemodynamic responses were then extracted from the three regions of interest (ROIs) that were used to define the optimal montage. Within each ROI, we extract the mean and standard deviation for fNIRS reconstructed signals across the vertices inside each ROI). (Figure 7.3.b). As a result, NIRSTORM allowed us to visualize fNIRS and EEG signals acquired simultaneously (Fig. 7.3.b). After considering 10 isolated bilateral bursts of polyspike and waves marked by an expert epileptologist, we estimated a significant negative fNIRS response at the sensor level. After applying cMEM reconstruction, the results suggested a left frontal fNIRS negative response with a clear decrease in HbO and a slight increase in HbR, peaking 19 seconds after the EEG discharge and returning to baseline 30 seconds after EEG discharge. This hemodynamic response could reflect either transient hypoxia or local inhibition associated with EEG interictal spikes^7^. It is worth mentioning that we did not find any significant hemodynamic response either in the homologous contralateral region (right frontal) or in the control region (left parietal).

## 7. Discussion

In this article, we introduced NIRSTORM, a fully integrated plugin implemented within Brainstorm environment. NIRSTORM is an easy-to-use and fully modular toolbox dedicated to fNIRS analysis ranging from experimental planning to 3D reconstruction along the cortical surface and analyses of multimodal signals.

### 7.1. fNIRS personalized optimal montage design

In section 3, we presented the concept of fNIRS optimal montage design and how it can be used to optimize the probe placement to target specific cortical regions, as further described in dedicated publications^14,15,31^. One of the main strengths and originality of this method is also its integration into Brainstorm. Indeed, Brainstorm provides user-friendly tools to define regions of interest along the cortical surface, which could then be considered as targeted regions for optimal montage design. The definition of these targeted ROIs could be defined quantitatively using preliminary fMRI results ^31^, EEG/MEG source imaging results^86^, anatomical atlases or drawn manually along the cortical surface.

Our proposed method is complementary to fNIRS optodes’ location decider^13^, which consists in choosing optodes from a set of predefined positions from the standard 10-10 or 10-05 international EEG systems, while estimating NIR light sensitivity maps on anatomical head templates, and Array designer ^16^, which ensures the estimation of a montage covering the full extent of the targeted region, but at the price of less density and overlap between fNIRS measurements. We acknowledge that one limitation of our current optimal montage algorithm lies in the spatial extent of the targeted region: the algorithm has been designed for the study of patients with focal epilepsy for whom the targeted region remains relatively small (20 to 60 cm^2^). Therefore, our optimization problem did not include any constraint on the coverage of the entire targeted region in the case of extended regions but rather focused more on providing a locally dense montage to improve the accuracy of local fNIRS 3D reconstruction. Ensuring a dense coverage of an extended region would necessarily require adding more sources and detectors. Whereas extended high-density fNIRS coverage has been demonstrated as essential to improve NIROT accuracy ^33^, we plan to implement strategies to combine constraints on channel density and coverage of more extended regions or networks in our future releases of NIRSTORM. Including these additional constraints would allow us to study the impact of the tradeoff between complete coverage of the targeted ROI and the spatial density of the montage on NIROT accuracy. It is worth mentioning that another popular tool to build fNIRS montage is AtlasViewer^28^ which is based on interactivity where the user will build the montage on a 2D grid and then project it on a subject head. This is especially useful when working with a cap and pre-designed grid, while estimating light sensitivity maps of the proposed montage. Such user-based interactivity makes it easy to modify the montage and assess resulting changes in the light sensitivity. In AtlasViewer, the estimation of montage sensitivity is also feasible using Monte Carlo simulation techniques (tMCimg^87^), either on the subject’s specific MRI or when using an anatomical template. As for our package, AtlasViewer also allows the use of an anatomical atlas to guide the definition of the targeted ROIs. However, the proposed approach remains mainly manual and does not guarantee that the obtained montage is optimal, and especially locally dense, as in our method proposed in NIRSTORM.

### 7.2. fNIRS channel space analysis

In section 4, we described how conventional channel space analysis of fNIRS data could be completed in NIRSTORM, including data importation, coregistration with anatomical data, preprocessing, estimation of the evoked hemodynamic response, and statistical analysis within a GLM framework. The integration with Brainstorm makes NIRSTORM user-friendly notably for the management of anatomical data and the coregistration between fNIRS sensors and the underlying anatomy. Additionally, every intermediate computation is saved in the database allowing for an easy visualization and quality control of every step of the proposed pipeline. To propose a user-friendly environment, we tried to to keep the number of NIRSTORM options for each processing step limited and only implement steps widely adopted in the community (for example, we decided to implement only two popular methods dedicated for fNIRS motion correction). However, this is not a limitation, since users can export their data in snirf data format^39^ at every step, and therefore apply any other pipeline of their choice outside NIRSTORM environment. Conversely, results from other fNIRS packages, when available in snirf data format, can also be imported back in NIRSTORM to benefit from Brainstorm/NIRSTORM advanced data visualization/interaction tools.

We acknowledge two limitations of the current GLM implementation in NIRSTORM: (i) our implementation assumes a fixed shape for the HRF model^55^, and (ii) the noise autocorrelation can be modeled using either precoloring or prewhitening with an auto-regressive process of order 1. Regarding the HRF, it has been shown in fMRI that its shape also carries relevant information^88^. We have also demonstrated that when considering locally dense optimal montage and NIROT reconstruction using MNE, we could accurately recover a variety of shapes of underlying HRF models through deconvolution methods^89^. We are planning to implement these HRF deconvolution approaches after NIROT reconstructions in future releases. Regarding noise autocorrelation, one particularity of NIRSTORM, in comparison to NIRS-SPM^22^, is that prewhitening is performed for each channel independently whereas NIRS-SPM assumes a uniform noise across channels. fNIRS noise characteristics and their implication on the GLM have been investigated in details in Huppert 2016^90^ and were implemented in the AnalyzIR Toolbox^27^. Their proposed robust GLM methods is entitled AR-IRLS (Autoregressive – Iteratively Reweighted Least Squares). This algorithm depends on two main concepts: (1) the noise is modeled as an auto-regressive process of order p, where p is estimated using a data-driven approach (2) ordinary least square algorithm is replaced by a robust regression using an iterative approach to weight down outliers and reduce the impact of motion^49,90,91^. Some of these advanced GLM approaches could be implemented in NIRSTORM in future releases, or data analyzed in other packages could also be imported back into NIRSTORM for multimodal visualization purposes.

### 7.3. fNIRS NIROT reconstruction along the cortical surface

In section 5, we introduced fNIRS analysis along the cortical surface, by presenting the estimation of fNIRS forward model using MCXLab toolbox^36^ before solving the NIROT inverse problem using either MNE or cMEM solvers. Regarding the forward model, NIRSTORM relies on the MCXlab toolbox, allowing accurate Monte Carlo simulation of the infrared photon transport within the head. It is important to note that the accuracy of NIR forward model depends on the accuracy of the segmentation of the anatomical MRI into five tissues (skin, skull, CSF, GM, and WM). NIRSTORM implementation offers a full integration of MCXlab tools as another Brainstorm plugin, therefore all these analyses can be completed within the same NIRSTORM environment, significantly facilitating their usage. It is worth mentioning that one limitation of this step might be the computation time required for the estimation of light fluences. This issue is becoming particularly relevant in the context of the optimal montage when large search spaces are considered since NIR light fluences should then be computed for every vertex inside the search space. To overcome this limitation, we also provide precomputed fluences on the anatomical template Colin27^17^, therefore allowing faster application of the optimal montage strategy on an anatomical template. The group managing MCXlab recently proposed a cloud service (https://mcx.space/cloud/) where users without GPU could compute fluences remotely. This service is available but has not yet been integrated into NIRSTORM. Other software packages were also proposed to solve the fNIRS forward problem. AtlasViewer^28^ is relying on tMCimg^87^ which is very similar to MCXLab. One limitation, however, is that tMCimg is not taking advantage of the massive parallelization power of GPU and reports that simulating a source with 100 million photons would take 6 hours on a 1 GHz Pentium 3 CPU (against a few minutes on a modern GPU). One common limitation of MCXLab and tMCimg is the use of voxel-based head tissue segmentations: using voxels to represent shapes with curved boundaries leads to expensive memory and computational costs because high voxel densities are usually needed to retain boundary accuracy. A better description of the head can however be obtained using arbitrary mesh such as unit volumes. This method is at the foundation of all the Finite Element Methods (FEM) and has the advantage of being able to create a fine description of the tissues near the boundaries (using small volumes elements) while allowing a coarse description inside homogenous tissues^92–94^. Such a method has notably been implemented in NIRfast^95^, which includes tools for optical modeling for small animal imaging, breast imaging, brain imaging, and light dose verification in photodynamic therapy of the pancreas. The impact of using FEM models in fNIRS was discussed in Tran et al 2020^93^ and extended in Yuan et al 2020^96^ showing that FEM could efficiently be used to model complex geometry, including complex vascular network geometry. It is also worth mentioning that several methods have been recently implemented in Brainstorm to generate high-quality mesh for the generation of the EEG/MEG forward model using FEM method (cf. DUNEuro^97^). fNIRS forward modeling could therefore also benefit from these FEM modeling tools. Indeed, it would be straightforward within Brainstorm to use such high-accuracy mesh and Mesh-Based Monte Carlo^93,96^ to generate the forward model for fNIRS. Moreover, this would allow the use of a unique and consistent anatomical representation of the head for EEG/MEG and fNIRS modeling.

Importantly, in section 6, we described several tools we implemented to probe the light sensitivity of our proposed fNIRS montages, featuring notably two measures of spatial overlap in fNIRS montage: (i) the minimum number of channels sensitive to a specific cortical region or (ii) the minimum sensitivity required to receive signals from that region from at least *N_c_* channels. We strongly believe that those tools are important to assess the coverage and density of fNIRS montages and should also be considered on more conventional sparse fNIRS montages that are not dedicated to NIROT 3D reconstruction. This approach is unique to NIRSTORM as other software such as AtlasViewer^28^ only allows the visualization of the normalized sensitivity in dB of the entire montage. One interesting feature of AtlasViewer is the estimation of localization error and resolution maps, which are key features when using fNIRS for 3D reconstruction as they provide quantitative measures of how accurate the location of an imaged activation centroid is, as well as the expected resolution. Implementing those maps using the resolution matrix is quite common in the context of EEG/MEG modeling^78,98,99^: it can be computed analytically for linear solutions such as MNE, whereas it can be estimated for nonlinear solvers such as cMEM^78^. We plan to include similar features to characterize the properties of NIROT in future releases of NIRSTORM.

One limitation of these tools is that light sensitivity values are not easily interpretable quantitatively, even though higher sensitivity is better and expected. Indeed, linking modeled light sensitivity values estimating using MCXlab with the actual amplitude of HbO/HbR measured fluctuations is not straightforward and depends on many other parameters such as probe installation, the fNIRS device to perform measurements, and recording conditions. One future use of photon simulation using MCXlab in NIRSTORM could be the estimation of subject-specific DPF/PVF, which would improve quantification performances. Indeed, in the current implementation, DPF values are set using an approximation proposed in Scholkmann et al, 2013^53^ and are independent of the region to be measured (although DPF has been shown to be region-dependent^100^). Zhao et al 2002^100^ reported that DPF estimations, made in frontal regions, might not hold for neonates or older adults. Since MCXlab allows us to simulate the photon trajectory inside the subject-specific head, it would be possible to estimate the distance traveled by the light in each tissue and use this estimate for the MBLL, making it specific to each individual^101^ (although the authors considered the head as a single-layer tissue).

Finally, one of the key features of NIRSTORM is the implementation of advanced NIROT models, i.e., MNE and cMEM solutions. cMEM solutions were implemented and validated in the Brain Entropy in Space and Time (BEst) plugin^67^, which is the core toolbox proposing MEM methods to solve EEG/MEG or fNIRS inverse problems. cMEM methodology has been evaluated for its ability to recover the spatial extent of the underlying generators from EEG/MEG signals^75,76,80,81,102^ and from fNIRS signals^31,35^. NIROT results using cMEM indeed suggest that cMEM was particularly well-suited for the reconstruction of HbO/HbR fluctuations on the cortical surface even in low SNR conditions. NIROT using cMEM was evaluated using realistic fNIRS simulations in Cai et al (2022)^35^ and when compared with fMRI during a finger tapping task^31^. cMEM was then successfully applied to assess fNIRS response to monitor changes in brain excitability elicited by transcranial magnetic stimulation^57^. The implementation of cMEM for NIROT reconstruction is a unique feature of NIRSTORM since most other fNIRS software such as AtlasViewer^28^ or NeuroDot^25,26^ proposing NIROT methods rely mainly on variants of MNE implementations. It is worth mentioning that cMEM reconstruction is performed in the time domain, where a nonlinear solution is estimated independently for every time point. We previously demonstrated that despite its nonlinear aspects, cMEM provides an accurate estimation of the underlying time courses of HbO and HbR responses, similar to the MNE solution (see Fig 7 and 8 and Cai et al 2021^31^). The nonlinear properties of cMEM lead to interesting spatial properties in NIROT maps, allowing an accurate estimation of the underlying spatial extent of the hemodynamic fluctuations, whereas MNE methods are not sensitive to this spatial extent (ref Cai et al Sc. Rep). Since we are dealing with slow hemodynamic signals, the time courses measured using fNIRS present specific regularities, especially temporal smoothness. These temporal regularities can also be exploited using a wavelet representation of the data representing the data in a time-frequency manner^103^. Wavelet time-frequency representation of fNIRS data has been considered appropriate for several applications such as the removal of physiological noise^104^, the estimation/deconvolution of the hemodynamic response at the sensors level^105^, and the reconstruction of fNIRS signals on the cortex^106^. In Abdelnour et al 2010^106^, the authors considered a time-frequency representation of fNIRS signals for NIROT. Still, they did not specifically evaluate the relevance of using wavelet representation when solving the NIROT inverse problem. We developed and validated a wavelet-based extension of MEM, denoted wMEM^107^, which is implemented in BEst toolbox in Brainstorm. wMEM has demonstrated interesting source imaging performances when localizing EEG/MEG oscillations during epileptic seizures^86^ or ongoing resting-state activity^82,102^. We are currently evaluating the accuracy of wMEM when reconstructing fNIRS slow oscillations during ongoing resting-state activity.

**Figure 8.**
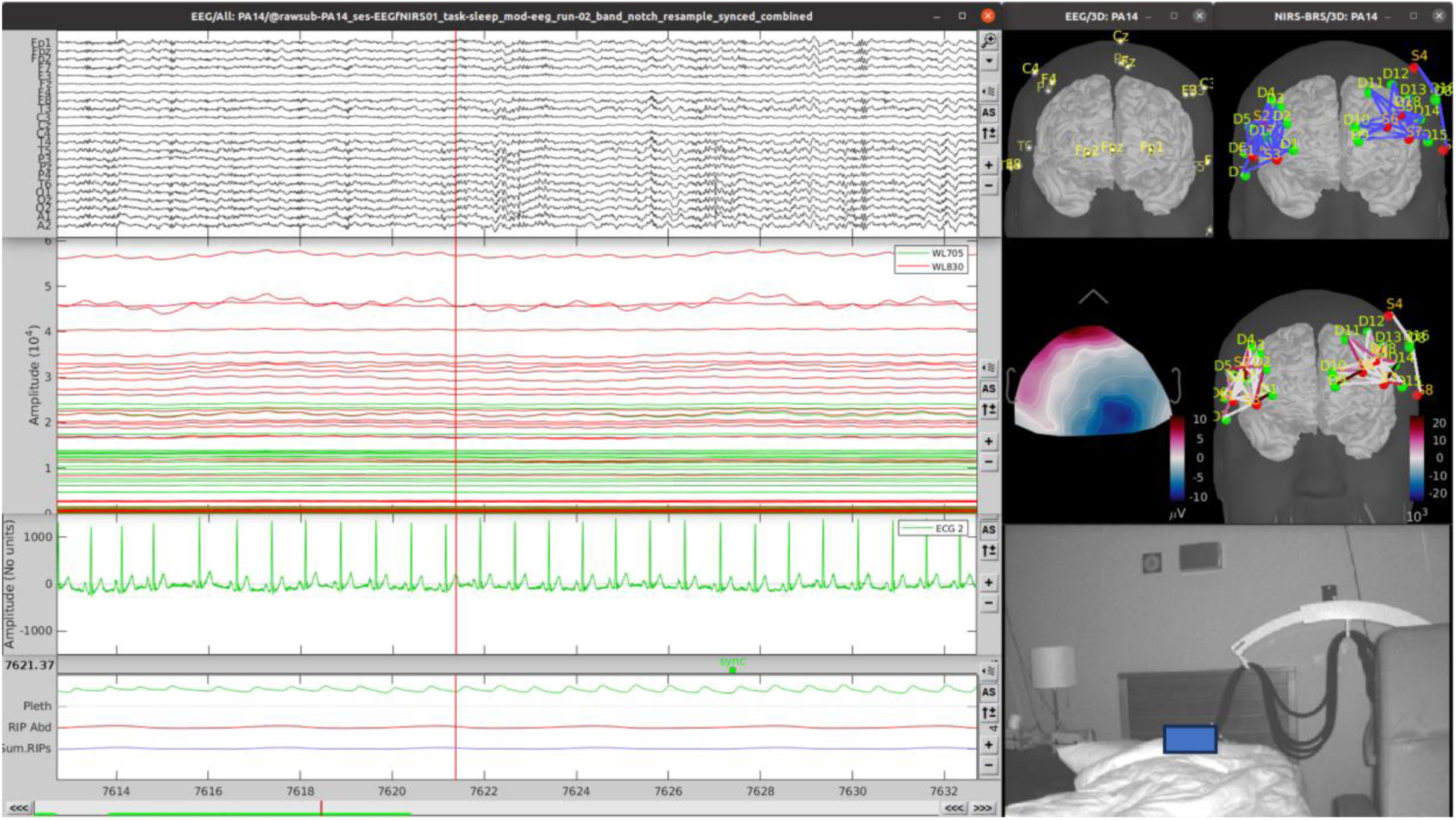
Multimodal data visualization. video-EEG-NIRS-polysomnography during sleep by showing on the left the recording of EEG, fNIRS, and additional measurements (ECG, pulse-oximetry (pleth), and respiration (RIP abd, sum.RIPs)). On the left, we can see the EEG and NIRS montage along with the topography of the signal for EEG and NIRS and the video.

Abdelnour et al, 2010^106^ proposed another Bayesian approach to solve the inverse problem, a method that has been implemented in the AnalyzIR toolbox^27^. In this study, the authors described how multiple priors could be integrated within the Restricted Maximum Likelihood framework to regularize the NIROT inverse problem. Interestingly, in this article, the anti-correlation between HbO and HbR signals is used to regularize the estimated time course on the cortical surface. Whereas NIRSTORM localizes each wavelength independently, they proposed to localize both wavelengths simultaneously, while adding prior on the anti-correlation between HbO and HbR. We believe that such prior could be relevant, especially in low SNR^108,109^ conditions where HbR time courses are often considered less reliable than HbO time courses. A similar approach could be considered to apply cMEM jointly on both wavelengths for NIROT reconstruction.

The same group also proposed a Bayesian fusion approach in the context of multimodal imaging fusion. Huppert et al 2008^110^ proposed a Bayesian fusion method dedicated to the joint reconstruction of simultaneous fNIRS and functional MRI data. They demonstrated that the high sampling rate of fNIRS data could be combined with the spatial information of MRI to generate spatiotemporal reconstructions of brain activity matching the benefits of both modalities. Importantly, they also proposed that the joint reconstruction of both modalities could cross-calibrate each modality therefore allowing the quantification of absolute micromolar HbO changes instead of relative measurements usually considered when using fNIRS or fMRI data alone. Similarly, Cao et al., 2021^111^ proposed a framework dedicated to joint EEG source localization and fNIRS NIROT reconstruction. They showed that using fNIRS localization as a spatial prior for EEG source localization could improve the spatial resolution of low-density EEG source imaging while preserving the excellent temporal resolution of EEG data. We believe that NIRSTORM/Brainstorm would therefore appear as an ideal multimodal environment to allow the future development of such multimodal fusion methods.

### 7.4. Multimodal integration

In section 6, we showed that NIRSTORM/Brainstorm could provide a comprehensive environment for the analysis of multimodal neurophysiological data. We presented how EEG/MEG source imaging could be considered to personalize fNIRS investigation, using the fNIRS optimal montage design. We then demonstrated how NIRSTORM/Brainstorm could be exploited to analyze EEG and fNIRS data acquired simultaneously, and how spontaneous epileptic discharges marked on scalp EEG data could be used to inform fNIRS analysis, allowing a constrained fusion approach. We then illustrated how NIROT using cMEM could provide promising 3D reconstruction maps of HbO/HbR responses elicited by epileptic discharges. Since both EEG and fNIRS signals can be imported in Brainstorm and synchronized in time, NIRSTORM/Brainstorm appears as an ideal environment to develop symmetrical data fusion approaches, where both modalities could be analyzed jointly based on a multimodal generative model^111,112^.

Regarding the inclusion of fMRI data within this multimodal framework, Brainstorm/NIRSTORM currently supports the importation of fMRI data or fMRI statistical results obtained from GLM analysis, for visualization in 3D or along the cortical surface, as illustrated in previous studies^31,57,75^. Therefore, fMRI data could be analyzed in other software^21,113–115^ before importing the results within Brainstorm. To interpolate fMRI volumetric results along the cortical surface that is usually considered for EEG/MEG source imaging or NIROT reconstructions, we implemented a Voronoi diagram interpolation scheme that is anatomically informed by the underlying anatomical sulco-gyral geometry^63^. Such a feature can be envisaged when considering previous fMRI findings to guide fNIRS optimal montage design^31^ or for multimodal fMRI/fNIRS comparisons. However, NIRSTORM currently proposes only limited support for joint analysis of fMRI and fNIRS signals. Fortunately, Brainstorm is an ideal multimodal environment that allows us to compare EEG/MEG source imaging and fMRI with intracranial EEG data^75^ or fNIRS NIROT results with the effects of transcranial magnetic stimulations^57^. Importantly, we have recently added in Brainstorm all the tools that are required to synchronize multiple recordings and visualize them together in a simple and user-friendly manner. Figure 8 shows that Brainstorm can effectively visualize simultaneous recording of video-EEG-NIRS-polysomnography during sleep by showing on the left the recording of EEG, fNIRS, and additional physiological measurements (ECG, pulse-oximetry, and respiration). Altogether, those tools allow for a comprehensive assessment of the neurophysiology and make Brainstorm a great tool for multimodal investigations.

### 7.5. Contributing to NIRSTORM development

NIRSTORM is an international team effort and welcomes any contribution by being an open-science initiative with its code hosted on GitHub under the GNU General Public License v3.0 (https://github.com/Nirstorm/nirstorm). Contributions can take multiple forms including questions in the Brainstorm forum, bug reports, tutorial writing, or the implantation of new functionalities. Between June 18^th^ 2021, and August 30^th^ 2024, NIRSTORM has been downloaded 562 times using the Brainstorm plugin system.

## 8. Conclusion

Over the past few years, NIRSTORM has grown to become a fully integrated plugin inside Brainstorm and is now able to perform most of the standard fNIRS analyses while benefitting from many key aspects of Brainstorm such as data structure, preprocessing, and visualization. Importantly, we have implemented two specific key features within NIRSTORM: (1) the optimal montage for personalized fNIRS, which enables one to efficiently optimize the fNIRS montage to target a specific cortical region of interest, and (2) an accurate and reliable NIROT reconstruction technique based on the Maximum Entropy on the Mean framework, which allows for accurate recovery of the spatial extent of hemodynamic fluctuations along the cortical surface. Additionally, NIRSTORM is a fully GUI-oriented software providing user-friendly visualization of fNIRS data and results, while allowing scripting features for reproducibility.

## 9. Appendix A – Description of cMEM

For NIROT we have adapted the “coherent” version of MEM, entitled cMEM, originally introduced in Chowdhury et al 2013^69^ and fully described in Chowdhury et al 2016^72^. The term “coherent” refers to the fact that we are using a coherent spatial prior, i.e. a data driven parcellation in *K* parcels stable over time, while the probability of being active evolves dynamically in time.

### The generative model of the data

The following linear model expresses the relationship between optical density fluctuations along the cortex ***j***(*t*) and fNIRS scalp measurement ***m***(*t*):

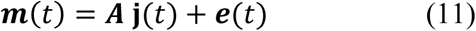

***m***(*t*) is the N_*C*_ –dimensional measurement vector for fNIRS signal (for a specific wavelength) at the time *t* where N_*C*_ denotes the number of fNIRS channels, ***j***(*t*) is the N_*v*_-dimensional vector denoting local fluctuations of optical density (for a specific wavelength) for N_*v*_ vertices of the cortical surface at time t. **A** is the ‘fNIRS forward model’ matrix, linking how local fluctuations of optical density at a specific vertex on the cortex are impacting channel data, with a dimension of N_*c*_ × N_*v*_. ***e***(*t*) models an additive measurement noise at time *t* measured on N_*C*_ channels. ***e***(*t*) is assumed to follow a normal distribution with zero mean and corresponding noise covariance structure **Σ**_**d**_ (N_*c*_ × N_*c*_ matrix).

The key feature of the cMEM Bayesian probabilistic framework is to consider a spatial prior model (or reference distribution) **v**, assuming that brain activity is organized within a set of *K* non-overlapping, and independent parcels. The activity of each parcel is then described independently from each other. Each parcel *k* composed of N_*k*_ vertices is characterized by an activation state S_*k*_, which is a hidden state variable controlling the activation of the parcel (the parcel being active when S_*k*_ = 1). The reference distribution ***v***_***k***_ for each parcel k is following a Bernoulli-Gaussian mixture distribution (Figure 8.c), sometimes also called a spike-and-slab prior, and defined as:

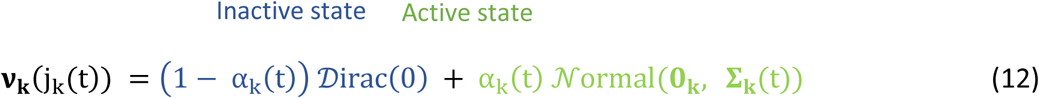

Where α_*k*_ is the probability that the *k^th^* parcel to be active (*Prob*(S_*k*_ = 1)). *Dirac* is a Dirac function which is used to switch off the parcel when the parcel is inactive (S_*k*_ = 0). *Normal*(**0**_**k**_, **Σ**_*k*_) denotes a Gaussian distribution of the optical densities of the *k^th^* parcel with mean **0**_**k**_ and spatial covariance matrix **Σ**_*k*_ describing the covariance of the N_*k*_ vertices within that *k^th^* parcel.

Since all the parcels are assumed to independent, this allows us to formulate the reference distribution for the whole brain **v** as the product of the distributions for each parcel.

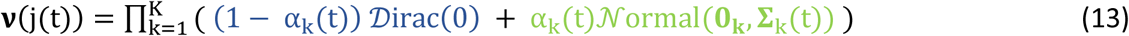

### Spatial parcellation

The spatial parcellation used in cMEM method is obtained using a data-driven parcellation technique^68^. To do so, whole brain parcellation was obtained using the Multiple Source Prelocalization (MSP)^68,116^ method which is a projection technique estimating a probability score assessing the contribution of every N_*v*_ vertex for its contribution to the scalp data. A stable spatial parcellation along the whole data window of interest could be obtained using a region-growing algorithm around best MSP scores while assuming a specific spatial neighborhood order along the cortical surface. By default, for NIROT, the spatial neighborhood parameter is tuned automatically so that the final number of parcels K is approximately equal to the number of fNIRS channels. (Figure 9.a)

**Figure 9.**
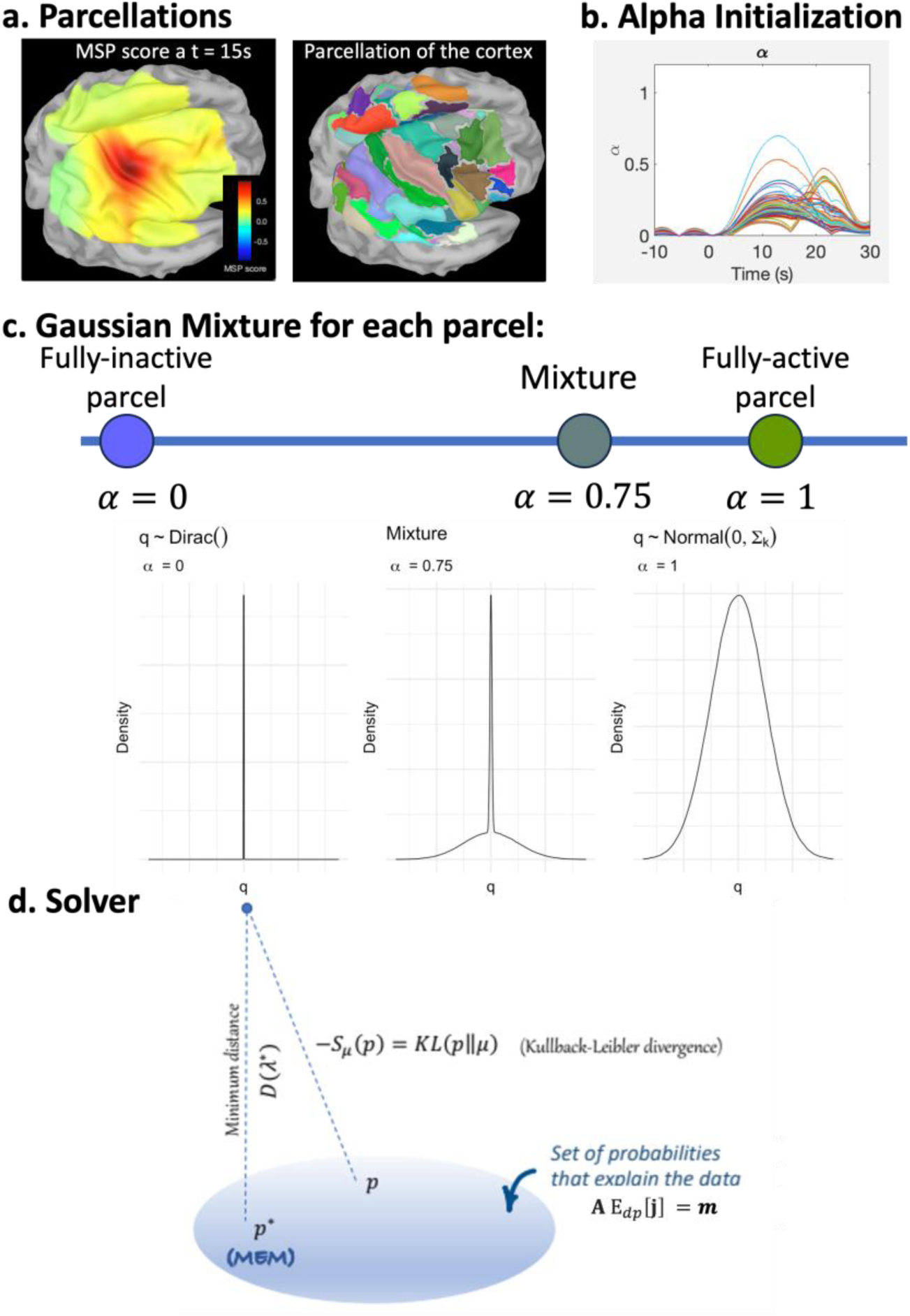
Theory of cMEM. a. Parcellation of the cortex in K non-overlapping parcels. b. Prior for the probability of each parcel being active (each line is a parcel). c. Prior for the activation of each parcel depending on their probability of being active. d. Solver. Showing the Kullback Leibler divergence between the prior and posterior distribution.

### Initialization of probability of hidden state variables

The probability of each parcel being active was initialized using the MSP scores used to create the parcellation. For each parcel *k*, α_*k*_ is estimated as the median MSP score within that parcel and evolves along time (see details in Appendix of Chowdhury et al 2013^69^). If an α_*k*_ reaches the threshold of 0.8, it is set to 1 automatically, meaning that the parcel is then considered fully active at that time point (See figure 9.b)

### Initialization of the spatial prior

The spatial covariance **Σ**_*k*_ of the Nornal distribution, when a parcel is active (see equation (13)), was initialized as follows:

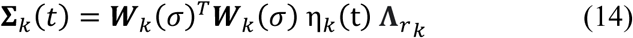

**Σ**_*k*_(*t*) is the optical densities covariance of parcel *k* at each time sample t, ***W***_*k*_(σ) is a spatial smoothness matrix which controls the local smoothness within the parcel using a diffusion model (to tune the level of spatial smoothness, σ is set to 0.6 following Chowdhury et al 2013^69^). 𝜂_*k*_(t) is defined as 5% of the energy of the depth weighted minimum norm estimate ***j***^^𝟐^ within the parcel *k* (see equation 15), and Λ_r*k*_ is the depth weighting matrix defined as the diagonal matrix of (**A**_*k*_^*T*^**A**_*k*_)^−ω^, where ***A***_*k*_ is the fNIRS forward model matrix for r_*k*_ vertices in parcel k.

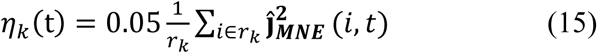

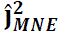 was computed as in section 5.3.1 considering the identity matrix for the noise covariance matrix Σ_𝑑_. When defining 𝜂_*k*_(t), the minimum norm solution was normalized by its maximum value for each time point, therefore allowing the mainly consider spatial information provided by the MNE solution only, rather than the amplitude (see details in Cai et al 2022 ^35^).

### The MEM Bayesian solver

Solving the MEM inverse problem consists in finding the probability distribution 𝑑𝑝 governing the fluctuation of optical densities along the cortex, the closest to our prior or reference distribution, and being able to explain the data on average. 𝑑𝑝^∗^will be estimated from the same probability family as our prior 𝑑𝜈. 𝑑𝑝^∗^ is found inside a set of probability dp so that:

1. 𝑑𝑝 is a probability density function with finite expectation
2. On average, 𝑑𝑝 is able to explain the measured data:

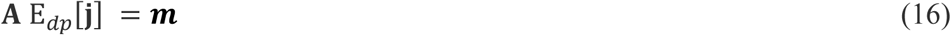

Where E_𝑑𝑝_[***j***] = ∫_ℝ_ ***j*** 𝑑𝑝(***j***) is the mathematical expectation of ***j*** with respect to the probability distribution 𝑑𝑝. We call C_m_ the set of all probability distributions on ***j*** that satisfies those two conditions. The notion of “distance” between probability distributions is defined using the Kullback-Leibler divergence 𝑆_ν_(𝑑𝑝) which defines the “distance” between any probability density function 𝑑𝑝 and our reference distribution **v**, with an associated probability density function 𝑑𝜈. This reference distribution is sometimes called the reference ‘null entropy’ since by construction 𝑆_ν_(𝑑𝜈) = 0.

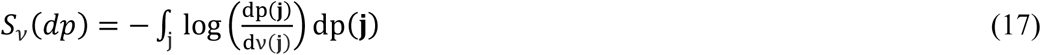

Importantly, 𝑆_ν_(𝑑𝑝), reported as the relative ‘Entropy drop’, measures the amount of information brought by the data with respect to our prior 𝑑𝜈. The MEM solution consists of finding the posterior distribution 𝑑𝑝^∗^that minimizes the Kullback-Leibler divergence (i.e. maximizes the relative entropy which is negative) while explaining the data on average (𝑑𝑝 ∈ *C*_𝑚_):

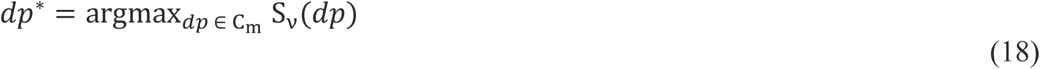

Finally, the MEM solution ***j***^∗^, the a posteriori mean estimate of the optical densities along the cortex is the mathematical expectation of the optical densities with respect to the optimal distribution 𝑑𝑝^∗^, i.e. ***j***^∗^ = E_𝑑𝑝_∗[***j***].

In practice, this constrained optimization problem is solved using its dual Legendre transform through the maximization of a concave function 𝐷(λ) defined in a space of dimension the number of channels (see proof in Amblard et al 2004^67^). cMEM solution is therefore obtained using the following 4 steps, which are repeated for every time sample *t* (see Figure 9.D):

1. Using the dual formulation of the MEM principle, solving the maximum entropy solution consists of maximizing the following function 𝐷(λ), which is a concave function of dimension the number of channels N_*c*_.

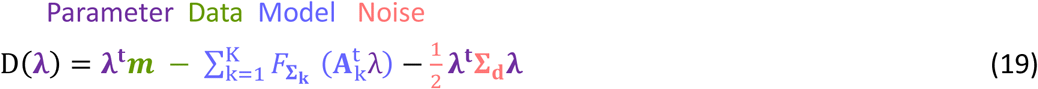 Where **A**_**k**_ is the forward problem matrix corresponding to the parcel ‘k’. Similarly, **Σ**_**k**_ corresponding to the sub-covariance matrix for the parcel ‘k’. 𝐹 is a function describing the free energy of the model inside a specific parcel k:

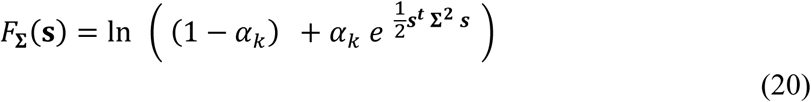 We can notice, that for inactive parcels, α_*k*_ = 0, the associated free energy 𝐹_**Σ**_(𝐬) is null (since the associated Dirac distribution has no free energy).
2. We find the set of parameters λ^∗^ that maximize 𝐷(λ) (i.e. minimize −𝐷(λ)). Importantly, it is important to notice that − 𝐷(λ) is a convex function and therefore has a unique minimum. Practically, λ^∗^(*t*) is estimated using a gradient descent algorithm for each time sample using fminfunc function in Matlab.
3. Once the optimal λ^∗^(*t*) is found, the final estimate of α_*k*_^∗^(*t*) for each parcel *k* at time *t* is estimated as follows (index *t* removed for simplicity):

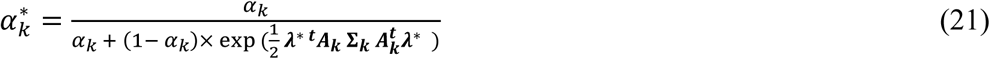
4. The final estimate of the amplitude of optical densities along the cortical surface inside each parcel k, for each time point *t*, i.e. the cMEM solution is obtained as follows

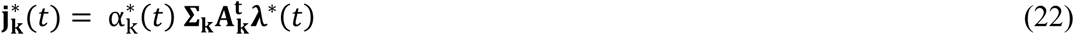

It is important here to realize that, like MNE, the cMEM solution is a linear function of λ, which is a vector of dimension of the number of channels N_*c*_. However, unlike in MNE, both α^∗^ and λ^∗^ have a non-linear relationship to the data, making cMEM a non-linear estimator.

## 10. Disclosure

The authors have no relevant financial or non-financial interests to disclose.

## 11. Acknowledgments/Funding Sources

This study was supported by Natural Sciences and Engineering Research Council of Canada (NSERC) Discovery grant, grant from Canadian Institutes of Health Research (CIHR) (PJT-159948), and the Fonds de Recherche du Québec—Nature et technologies (FRQNT) Research team grants held by CG. fNIRS equipment was acquired using grants from NSERC Research Tools and Instrumentation Program and the Canadian Foundation for Innovation (CG). From 2018 to 2023, FT 2018-2023 was supported by an NIH grant for USC: “Signal & Image Processing Institute, University of Southern California, Los Angeles, CA USA”

## 12. Code and Data Availability

The code of NIRSTORM is currently hosted on GitHub https://github.com/Nirstorm/nirstorm under the license GNU General Public License v3.0. Tapping Data used in this tutorial can be accessed at https://osf.io/md54y/?view_only=0d8ad17d1e1449b5ad36864eeb3424ed.

A tutorial to reproduce the figure can be found at https://neuroimage.usc.edu/brainstorm/Tutorials/NIRSTORM

## Notes

### Competing Interest Statement

The authors have declared no competing interest.

## Reference

1. Tak S, Ye JC. Statistical analysis of fNIRS data: a comprehensive review. Neuroimage. 2014;85:72–91. doi:10.1016/j.neuroimage.2013.06.016

2. Hoshi Y. Hemodynamic Signals in FNIRS. Vol 225. 1st ed. Elsevier B.V.; 2016. doi:10.1016/bs.pbr.2016.03.004

3. Yücel MA, Selb JJ, Huppert T, Franceschini MA, Boas DA. Functional Near Infrared Spectroscopy: Enabling routine functional brain imaging. Curr Opin Biomed Eng. 2017;4:78–86. doi:10.1016/j.cobme.2017.09.011

4. Scholkmann F, Kleiser S, Metz AJ, et al. A review on continuous wave functional near-infrared spectroscopy and imaging instrumentation and methodology. Neuroimage. 2014;85:6–27. doi:10.1016/j.neuroimage.2013.05.004

5. Talamonti D, Vincent T, Fraser S, Nigam A, Lesage F, Bherer L. The benefits of physical activity in individuals with cardiovascular risk factors: A longitudinal investigation using fNIRS and dual-task walking. J Clin Med. 2021;10(4):1–14. doi:10.3390/jcm10040579

6. Ren H, Jiang X, Xu K, et al. A Review of Cerebral Hemodynamics During Sleep Using Near-Infrared Spectroscopy. Front Neurol. 2020;11(November):1–19. doi:10.3389/fneur.2020.524009

7. Pellegrino G, Machado A, von Ellenrieder N, et al. Hemodynamic Response to Interictal Epileptiform Discharges Addressed by Personalized EEG-fNIRS Recordings. Front Neurosci. 2016;10:102. doi:10.3389/fnins.2016.00102

8. Kassab A, Hinnoutondji Toffa D, Robert M, Lesage F, Peng K, Khoa Nguyen D. Hemodynamic changes associated with common EEG patterns in critically ill patients: Pilot results from continuous EEG-fNIRS study. Neuroimage Clin. 2021;32. doi:10.1016/j.nicl.2021.102880

9. Buxton RB. Introduction to Functional Magnetic Resonance Imaging: Principles and Techniques. *null*. Published online 2002. doi: null

10. Boas DA, Dale AM, Franceschini MA. Diffuse optical imaging of brain activation: Approaches to optimizing image sensitivity, resolution, and accuracy. Neuroimage. 2004;23(SUPPL. 1). doi:10.1016/j.neuroimage.2004.07.011

11. Gagnon L, Yücel MA, Dehaes M, et al. Quantification of the cortical contribution to the NIRS signal over the motor cortex using concurrent NIRS-fMRI measurements. Neuroimage. 2012;59(4):3933–3940. doi:10.1016/j.neuroimage.2011.10.054

12. Yücel MA, Lühmann A v., Scholkmann F, et al. Best practices for fNIRS publications. Neurophotonics. 2021;8(01):1–34. doi:10.1117/1.nph.8.1.012101

13. Zimeo Morais GA, Balardin JB, Sato JR. FNIRS Optodes’ Location Decider (fOLD): A toolbox for probe arrangement guided by brain regions-of-interest. Sci Rep. 2018;8(1):1–11. doi:10.1038/s41598-018-21716-z

14. Machado A, Marcotte O, Lina JM, Kobayashi E, Grova C. Optimal optode montage on electroencephalography/functional near-infrared spectroscopy caps dedicated to study epileptic discharges. J Biomed Opt. 2014;19(2):026010. doi:10.1117/1.jbo.19.2.026010

15. Machado A, Cai Z, Pellegrino G, et al. Optimal positioning of optodes on the scalp for personalized functional near-infrared spectroscopy investigations. J Neurosci Methods. 2018;309(August):91–108. doi:10.1016/j.jneumeth.2018.08.006

16. Brigadoi S, Salvagnin D, Fischetti M, Cooper RJ. Array Designer: automated optimized array design for functional near-infrared spectroscopy. Neurophotonics. Published online 2018. doi:10.1117/1.nph.5.3.035010

17. Holmes CJ, Hoge R, Collins L, Woods R, Toga AW, Evans AC. Enhancement of MR images using registration for signal averaging. J Comput Assist Tomogr. 1998;22(2). doi:10.1097/00004728-199803000-00032

18. Fonov V, Evans A, McKinstry R, Almli C, Collins D. Unbiased nonlinear average age-appropriate brain templates from birth to adulthood. Neuroimage. 2009;47. doi:10.1016/s1053-8119(09)70884-5

19. Gramfort A, Luessi M, Larson E, et al. MEG and EEG data analysis with MNE-Python. Front Neurosci. 2013;(7 DEC). doi:10.3389/fnins.2013.00267

20. Oostenveld R, Fries P, Maris E, Schoffelen JM. FieldTrip: Open source software for advanced analysis of MEG, EEG, and invasive electrophysiological data. Comput Intell Neurosci. 2011;2011. doi:10.1155/2011/156869

21. Henson RN, Abdulrahman H, Flandin G, Litvak V. Multimodal integration of M/EEG and f/MRI data in SPM12. Front Neurosci. 2019;13(APR). doi:10.3389/fnins.2019.00300

22. Ye JC, Tak S, Jang KE, Jung J, Jang J. NIRS-SPM: Statistical parametric mapping for near-infrared spectroscopy. Neuroimage. 2009;44(2). doi:10.1016/j.neuroimage.2008.08.036

23. Tremblay J, Martínez-Montes E, Hüsser A, et al. LIONirs: Flexible Matlab toolbox for fNIRS data analysis. bioRxiv. Published online 2020:1–25. doi:10.1101/2020.09.11.257634

24. Hou X, Zhang Z, Zhao C, et al. NIRS-KIT: a MATLAB toolbox for both resting-state and task fNIRS data analysis. Neurophotonics. 2021;8(01). doi:10.1117/1.nph.8.1.010802

25. Speh E, Thacker Y, Segel A, et al. NeuroDOT: A Python Neuroimaging Toolbox for DOT. In: Optics InfoBase Conference Papers.; 2022. doi:10.1364/boda.2023.jtu4b.24

26. Eggebrecht AT, Culver JP. Neurodot: An extensible Matlab toolbox for streamlined optical functional mapping. In: Optics InfoBase Conference Papers. Vol Part F142-ECBO 2019.; 2019. doi:10.1117/12.2527164

27. Santosa H, Zhai X, Fishburn F, Huppert T. The NIRS brain AnalyzIR toolbox. Algorithms. 2018;11(5):73. doi:10.3390/a11050073

28. Aasted CM, Yücel MA, Cooper RJ, et al. Anatomical guidance for functional near-infrared spectroscopy: AtlasViewer tutorial. Neurophotonics. 2015;2(2). doi:10.1117/1.nph.2.2.020801

29. Tadel F, Bock E, Niso G, et al. MEG/EEG group analysis with brainstorm. Front Neurosci. 2019;13(FEB):1–21. doi:10.3389/fnins.2019.00076

30. Tadel F, Baillet S, Mosher JC, Pantazis D, Leahy RM. Brainstorm: a user-friendly application for MEG/EEG analysis. Comput Intell Neurosci. 2011;2011:8.

31. Cai Z, Uji M, Aydin Ü, et al. Evaluation of a personalized functional near infra-red optical tomography workflow using maximum entropy on the mean. Hum Brain Mapp. 2021;42(15):4823–4843. doi:10.1002/hbm.25566

32. Manual CU. Ibm ilog cplex optimization studio. Version. 1987;12(1987-2018):1.

33. Wheelock MD, Culver JP, Eggebrecht AT. High-density diffuse optical tomography for imaging human brain function. Review of Scientific Instruments. 2019;90(5). doi:10.1063/1.5086809

34. Machado A, Cai Z, Pellegrino G, et al. Optimal positioning of optodes on the scalp for personalized functional near-infrared spectroscopy investigations. J Neurosci Methods. 2018;309:91–108. doi:10.1016/j.jneumeth.2018.08.006

35. Cai Z, Machado A, Chowdhury RA, et al. Diffuse optical reconstructions of functional near infrared spectroscopy data using maximum entropy on the mean. Sci Rep. 2022;12(1):1–18. doi:10.1038/s41598-022-06082-1

36. Fang Q, Boas DA. Monte Carlo Simulation of Photon Migration in 3D Turbid Media Accelerated by Graphics Processing Units. Opt Express. 2009;17(22):20178. doi:10.1364/oe.17.020178

37. Land AH, Doig AG. An Automatic Method of Solving Discrete Programming Problems. Econometrica. 1960;28(3). doi:10.2307/1910129

37. Huppert T, Diamond SG, Franceschini MA, Boas DA. HomER: a review of time-series analysis methods for near-infrared spectroscopy of the brain. Appl Opt. 2009;48:D280-98.

39. Tucker S, Dubb J, Kura S, et al. Introduction to the shared near infrared spectroscopy format. Neurophotonics. 2022;10(01). doi:10.1117/1.nph.10.1.013507

40. Gaser C, Dahnke R, Kurth F LE. CAT – A Computational Anatomy Toolbox for the Analysis of Structural MRI Data. NeuroImage, in review.

41. Dale A, Fischl B, Sereno MI. Cortical Surface-Based Analysis: I. Segmentation and Surface Reconstruction. Neuroimage. 1999;9(2):179–194.

42. Pollonini L, Bortfeld H, Oghalai JS. PHOEBE: a method for real time mapping of optodes-scalp coupling in functional near-infrared spectroscopy. Biomed Opt Express. 2016;7(12):5104. doi:10.1364/boe.7.005104

43. Brown CE. Applied Multivariate Statistics in Geohydrology and Related Sciences. Technometrics. 1998;43(1):110–110. doi:10.1198/tech.2001.s566

44. Cooper RJ, Selb J, Gagnon L, et al. A systematic comparison of motion artifact correction techniques for functional near-infrared spectroscopy. Front Neurosci. 2012;6(OCT):1–10. doi:10.3389/fnins.2012.00147

45. Scholkmann F, Spichtig S, Muehlemann T, Wolf M. How to detect and reduce movement artifacts in near-infrared imaging using moving standard deviation and spline interpolation. Physiol Meas. 2010;31(5):649–662. doi:10.1088/0967-3334/31/5/004

46. Fishburn FA, Ludlum RS, Vaidya CJ, Medvedev A V. Temporal Derivative Distribution Repair (TDDR): A motion correction method for fNIRS. Neuroimage. 2019;184(September 2018):171–179. doi:10.1016/j.neuroimage.2018.09.025

47. Scholkmann F, Tachtsidis I, Wolf M, Wolf U. Systemic physiology augmented functional near-infrared spectroscopy: a powerful approach to study the embodied human brain. Neurophotonics. 2022;9(3):030801. doi:10.1117/1.NPh.9.3.030801

48. Wyser D, Mattille M, Wolf M, Lambercy O, Scholkmann F, Gassert R. Short-channel regression in functional near-infrared spectroscopy is more effective when considering heterogeneous scalp hemodynamics. Neurophotonics. 2020;7(03). doi:10.1117/1.nph.7.3.035011

49. Santosa H, Zhai X, Fishburn F, Sparto PJ, Huppert TJ. Quantitative comparison of correction techniques for removing systemic physiological signal in functional near-infrared spectroscopy studies. Neurophotonics. 2020;7(03):1–21. doi:10.1117/1.nph.7.3.035009

50. Gagnon L, Perdue K, Greve DN, Goldenholz D, Kaskhedikar G, Boas DA. Improved recovery of the hemodynamic response in diffuse optical imaging using short optode separations and state-space modeling. Neuroimage. 2011;56(3). doi:10.1016/j.neuroimage.2011.03.001

51. von Lühmann A, Li X, Müller KR, Boas DA, Yücel MA. Improved physiological noise regression in fNIRS: A multimodal extension of the General Linear Model using temporally embedded Canonical Correlation Analysis. Neuroimage. 2020;208(September 2019):116472. doi:10.1016/j.neuroimage.2019.116472

52. Duncan A, Meek JH, Clemence M, et al. Measurement of Cranial Optical Path Length as a Function of Age Using Phase Resolved Near Infrared Spectroscopy. Pediatr Res. 1996;39(5):889–894. doi:10.1203/00006450-199605000-00025

53. Scholkmann F, Wolf M. General equation for the differential pathlength factor of the frontal human head depending on wavelength and age. J Biomed Opt. 2013;18(10):105004. doi:10.1117/1.jbo.18.10.105004

54. Monti M. Statistical analysis of fMRI time-series: A critical review of the GLM approach. Front Hum Neurosci. 2011;5(March):1–13. doi:10.3389/fnhum.2011.00028

55. Glover GH. Deconvolution of impulse response in event-related BOLD fMRI. Neuroimage. Published online 1999. doi:10.1006/nimg.1998.0419

56. Huppert TJ, Allen MS, Diamond SG, Boas DA. Estimating cerebral oxygen metabolism from fMRI with a dynamic multicompartment windkessel model. Hum Brain Mapp. 2009;30(5):1548–1567. doi:10.1002/hbm.20628

57. Cai Z, Pellegrino G, Lina JM, Benali H, Grova C. Hierarchical Bayesian modeling of the relationship between task-related hemodynamic responses and cortical excitability. Hum Brain Mapp. 2022;(June):1–25. doi:10.1002/hbm.26107

58. Tak S, Ye JC. Statistical analysis of fNIRS data: a comprehensive review. Neuroimage. 2014;85:72–91. doi:10.1016/j.neuroimage.2013.06.016

59. Benjamini Y, Hochberg Y. Controlling the False Discovery Rate: A Practical and Powerful Approach to Multiple Testing. Journal of the Royal Statistical Society: Series B (Methodological). 1995;57(1):289–300. doi:10.1111/j.2517-6161.1995.tb02031.x

60. Calvetti D, Somersalo E. Inverse problems: From regularization to Bayesian inference. Wiley Interdiscip Rev Comput Stat. 2018;10(3):1–19. doi:10.1002/wics.1427

61. Arridge SR. Optical tomography in medical imaging. Inverse Probl. 1999;15(2). doi:10.1088/0266-5611/15/2/022

62. Grech R, Cassar T, Muscat J, et al. Review on solving the inverse problem in EEG source analysis. J Neuroeng Rehabil. 2008;5:1–33. doi:10.1186/1743-0003-5-25

63. Grova C, Makni S, Flandin G, Ciuciu P, Gotman J, Poline JB. Anatomically informed interpolation of fMRI data on the cortical surface. Neuroimage. 2006;31(4):1475–1486. doi:10.1016/j.neuroimage.2006.02.049

64. Lin FH, Witzel T, Ahlfors SP, Stufflebeam SM, Belliveau JW, Hämäläinen MS. Assessing and improving the spatial accuracy in MEG source localization by depth-weighted minimum-norm estimates. Neuroimage. 2006;31(1). doi:10.1016/j.neuroimage.2005.11.054

65. Samuelsson JG, Peled N, Mamashli F, Ahveninen J, Hämäläinen MS. Spatial fidelity of MEG/EEG source estimates: A general evaluation approach. Neuroimage. 2021;224(October 2020):117430. doi:10.1016/j.neuroimage.2020.117430

66. Hämäläinen MS, Ilmoniemi RJ. Interpreting magnetic fields of the brain: minimum norm estimates. Med Biol Eng Comput. 1994;32(1). doi:10.1007/BF02512476

67. Amblard C, Lapalme E, Lina JM. Biomagnetic Source Detection by Maximum Entropy and Graphical Models. IEEE Trans Biomed Eng. 2004;51(3):427–442. doi:10.1109/TBME.2003.820999

68. Lapalme E, Lina JM, Mattout J. Data-driven parceling and entropic inference in MEG. Neuroimage. 2006;30(1):160–171. doi:10.1016/j.neuroimage.2005.08.067

69. Chowdhury RA, Lina JM, Kobayashi E, Grova C. MEG Source Localization of Spatially Extended Generators of Epileptic Activity: Comparing Entropic and Hierarchical Bayesian Approaches. PLoS One. 2013;8(2). doi:10.1371/journal.pone.0055969

70. Hansen PC. The L-Curve and its Use in the Numerical Treatment of Inverse Problems. In: In Computational Inverse Problems in Electrocardiology, Ed. P. Johnston, Advances in Computational Bioengineering. WIT Press; 2000:119–142.

71. Grova C, Daunizeau J, Lina JM, Bénar CG, Benali H, Gotman J. Evaluation of EEG localization methods using realistic simulations of interictal spikes. Neuroimage. 2006;29(3). doi:10.1016/j.neuroimage.2005.08.053

72. Chowdhury RA, Merlet I, Birot G, et al. Complex patterns of spatially extended generators of epileptic activity: Comparison of source localization methods cMEM and 4-ExSo-MUSIC on high resolution EEG and MEG data. Neuroimage. 2016;143:175–195. doi:10.1016/j.neuroimage.2016.08.044

73. Heers M, Hedrich T, An D, et al. Spatial correlation of hemodynamic changes related to interictal epileptic discharges with electric and magnetic source imaging. Hum Brain Mapp. 2014;35(9):4396–4414. doi:10.1002/hbm.22482

74. Grova C, Aiguabella M, Zelmann R, Lina JM, Hall JA, Kobayashi E. Intracranial EEG potentials estimated from MEG sources: A new approach to correlate MEG and iEEG data in epilepsy. Hum Brain Mapp. 2016;37(5). doi:10.1002/hbm.23127

75. Abdallah C, Hedrich T, Koupparis A, et al. Clinical Yield of Electromagnetic Source Imaging and Hemodynamic Responses in Epilepsy: Validation With Intracerebral Data. Neurology. Published online 2022.

76. Pellegrino G, Hedrich T, Chowdhury RA, et al. Clinical yield of magnetoencephalography distributed source imaging in epilepsy: A comparison with equivalent current dipole method. Hum Brain Mapp. 2018;39(1). doi:10.1002/hbm.23837

77. Pellegrino G, Hedrich T, Porras-Bettancourt M, et al. Accuracy and spatial properties of distributed magnetic source imaging techniques in the investigation of focal epilepsy patients. Hum Brain Mapp. 2020;41(11):3019–3033. doi:10.1002/hbm.24994

78. Hedrich T, Pellegrino G, Kobayashi E, Lina JM, Grova C. Comparison of the spatial resolution of source imaging techniques in high-density EEG and MEG. Neuroimage. 2017;157(June):531–544. doi:10.1016/j.neuroimage.2017.06.022

79. Chowdhury RA, Pellegrino G, Aydin Ü, et al. Reproducibility of EEG-MEG fusion source analysis of interictal spikes: Relevance in presurgical evaluation of epilepsy. Hum Brain Mapp. 2018;39(2). doi:10.1002/hbm.23889

80. Avigdor T, Abdallah C, Afnan J, et al. Consistency of electrical source imaging in presurgical evaluation of epilepsy across different vigilance states. Ann Clin Transl Neurol. 2024;11(2). doi:10.1002/acn3.51959

81. Afnan J, Cai Z, Lina JM, et al. EEG/MEG source imaging of deep brain activity within the maximum entropy on the mean framework: Simulations and validation in epilepsy. Hum Brain Mapp. 2024;45(10):e26720. 10.1002/hbm.26720

82. Aydin Ü, Pellegrino G, Ali OBK b., et al. Magnetoencephalography resting state connectivity patterns as indicatives of surgical outcome in epilepsy patients. J Neural Eng. 2020;17(3). doi:10.1088/1741-2552/ab8113

83. Chen WL, Wagner J, Heugel N, et al. Functional Near-Infrared Spectroscopy and Its Clinical Application in the Field of Neuroscience: Advances and Future Directions. Front Neurosci. 2020;14(July):1–15. doi:10.3389/fnins.2020.00724

84. Cai Z, Pellegrino G, Spilkin A, et al. Hemodynamic correlates of fluctuations in neuronal excitability: A simultaneous Paired Associative Stimulation (PAS) and functional near infra-red spectroscopy (fNIRS) study. Neuroimage: Reports. 2022;2(3):100099. doi:10.1016/j.ynirp.2022.100099

85. Schneider W, Eschman A, Zuccolotto A. E-prime computer software and manual. Pittsburgh, PA: Psychology Software Tools *…*. Published online 2002.

86. Pellegrino G, Hedrich T, Chowdhury R, et al. Source localization of the seizure onset zone from ictal EEG/MEG data. Hum Brain Mapp. 2016;37(7):2528–2546. doi:10.1002/hbm.23191

87. Boas DA, Culver JP, Stott JJ, Dunn AK. Three dimensional Monte Carlo code for photon migration through complex heterogeneous media including the adult human head. Opt Express. 2002;10(3). doi:10.1364/oe.10.000159

88. Chen G, Taylor PA, Reynolds RC, et al. BOLD Response is more than just magnitude: Improving detection sensitivity through capturing hemodynamic profiles. Neuroimage. 2023;277. doi:10.1016/j.neuroimage.2023.120224

89. Machado A, Cai Z, Vincent T, et al. Deconvolution of hemodynamic responses along the cortical surface using personalized functional near infrared spectroscopy. Sci Rep. 2021;11(1). doi:10.1038/s41598-021-85386-0

90. Huppert TJ. Commentary on the statistical properties of noise and its implication on general linear models in functional near-infrared spectroscopy. Neurophotonics. 2016;3(1):010401. doi:10.1117/1.nph.3.1.010401

91. Barker JW, Aarabi A, Huppert TJ. Autoregressive model based algorithm for correcting motion and serially correlated errors in fNIRS. Biomed Opt Express. 2013;4(8):1366–1379. doi:10.1364/boe.4.001366

92. Fang Q, Boas DA. Tetrahedral mesh generation from volumetric binary and grayscale images. In: Proceedings – 2009 IEEE International Symposium on Biomedical Imaging: From Nano to Macro, ISBI 2009.; 2009. doi:10.1109/ISBI.2009.5193259

93. Tran AP, Yan S, Fang Q. Improving model-based functional near-infrared spectroscopy analysis using mesh-based anatomical and light-transport models. Neurophotonics. 2020;7(01). doi:10.1117/1.nph.7.1.015008

94. Jermyn M, Ghadyani H, Mastanduno MA, et al. Fast segmentation and high-quality three-dimensional volume mesh creation from medical images for diffuse optical tomography. J Biomed Opt. 2013;18(8). doi:10.1117/1.jbo.18.8.086007

95. Dehghani H, Eames ME, Yalavarthy PK, et al. Near infrared optical tomography using NIRFAST: Algorithm for numerical model and image reconstruction. Commun Numer Methods Eng. 2009;25(6). doi:10.1002/cnm.1162

96. Yuan Y, Yan S, Fang Q. Light transport modeling in highly complex tissues using implicit mesh-based Monte Carlo algorithm. Biomed Opt Express. Published online November 25, 2020. doi:10.1364/boe.411898

97. Medani T, Garcia-Prieto J, Tadel F, et al. Brainstorm-DUNEuro: An integrated and user-friendly Finite Element Method for modeling electromagnetic brain activity. Neuroimage. 2023;267. doi:10.1016/j.neuroimage.2022.119851

98. Schoffelen JM, Gross J. Source connectivity analysis with MEG and EEG. Hum Brain Mapp. 2009;30(6). doi:10.1002/hbm.20745

99. Grave De Peralta Menendez R, Hauk O, Andino SG, Vogt H, Michel C. Linear inverse solutions with optimal resolution kernels applied to electromagnetic tomography. Hum Brain Mapp. 1997;5(6). doi:10.1002/(SICI)1097-0193(1997)5:6<454::AID-HBM6>3.0.CO;2-2

100. Zhao H, Tanikawa Y, Gao F, et al. Maps of optical differential pathlength factor of human adult forehead, somatosensory motor and occipital regions at multi-wavelengths in NIR. Phys Med Biol. 2002;47(12). doi:10.1088/0031-9155/47/12/306

101. Chatterjee S, Phillips JP, Kyriacou PA. Differential pathlength factor estimation for brain-like tissue from a single-layer Monte Carlo model. In: *Proceedings of the Annual International Conference of the IEEE Engineering in Medicine and Biology Society*, EMBS. Vol 2015-November.; 2015. doi:10.1109/EMBC.2015.7319092

102. Afnan J, von Ellenrieder N, Lina JM, et al. Validating MEG source imaging of resting state oscillatory patterns with an intracranial EEG atlas. Neuroimage. Published online May 2023:120158. doi:10.1016/j.neuroimage.2023.120158

103. Lina JM, Mayrand M. Complex Daubechies Wavelets. Appl Comput Harmon Anal. 1995;2(3):219–229. doi:10.1006/acha.1995.1015

104. Duan L, Zhao Z, Lin Y, Wu X, Luo Y, Xu P. Wavelet-based method for removing global physiological noise in functional near-infrared spectroscopy. Biomed Opt Express. 2018;9(8):3805. doi:10.1364/boe.9.003805

105. Lina JM, Matteau-Pelletier C, Dehaes M, Desjardins M, Lesage F. Wavelet-based estimation of the hemodynamic responses in diffuse optical imaging. Med Image Anal. 2010;14(4):606–616. doi:10.1016/j.media.2010.04.006

106. Abdelnour F, Genovese C, Huppert T. Hierarchical Bayesian regularization of reconstructions for diffuse optical tomography using multiple priors. Biomed Opt Express. 2010;1(4):1084. doi:10.1364/BOE.1.001084

107. Lina JM, Chowdhury R, Lemay E, Kobayashi E, Grova C. Wavelet-based localization of oscillatory sources from magnetoencephalography data. IEEE Trans Biomed Eng. 2014;61(8). doi:10.1109/TBME.2012.2189883

108. Culver JP, Siegel AM, Franceschini MA, Mandeville JB, Boas DA. Evidence that cerebral blood volume can provide brain activation maps with better spatial resolution than deoxygenated hemoglobin. Neuroimage. 2005;27(4). doi:10.1016/j.neuroimage.2005.05.052

109. Shader MJ, Gramfort A, Larson E. Oxygenated hemoglobin signal provides greater predictive performance of experimental condition than de-oxygenated. bioRxiv Neuroscience. Published online 2021. https://www.biorxiv.org/content/10.1101/2021.11.19.469225v1?rss=1&utm_source=researcher_app&utm_medium=referral&utm_campaign=RESR_MRKT_Researcher_inbound

110. Huppert TJ, Diamond SG, Boas DA. Direct estimation of evoked hemoglobin changes by multimodality fusion imaging. J Biomed Opt. 2008;13(5). doi:10.1117/1.2976432

111. Cao J, Huppert TJ, Grover P, Kainerstorfer JM. Enhanced spatiotemporal resolution imaging of neuronal activity using joint electroencephalography and diffuse optical tomography. Neurophotonics. 2021;8(01):015002. doi:10.1117/1.NPh.8.1.015002

112. Daunizeau J, Grova C, Marrelec G, et al. Symmetrical event-related EEG/fMRI information fusion in a variational Bayesian framework. Neuroimage. 2007;36(1):69–87. doi:10.1016/j.neuroimage.2007.01.044

113. Esteban O, Markiewicz CJ, Blair RW, et al. fMRIPrep: a robust preprocessing pipeline for functional MRI. Nat Methods. 2019;16(1):111–116. doi:10.1038/s41592-018-0235-4

114. Woolrich MW, Jbabdi S, Patenaude B, et al. Bayesian analysis of neuroimaging data in FSL. Neuroimage. 2009;45(1 Suppl):S173–S186. doi:10.1016/j.neuroimage.2008.10.055

115. Cox RW. AFNI: What a long strange trip it’s been. Neuroimage. 2012;62(2). doi:10.1016/j.neuroimage.2011.08.056

116. Mattout J, Pélégrini-issac M, Garnero L, Benali H. Multivariate source prelocalization (MSP): Use of functionally informed basis functions for better conditioning the MEG inverse problem. Neuroimage. 2005;26(2):356–373. doi:10.1016/j.neuroimage.2005.01.026

